# Epigenetic mechanisms of partial dosage compensation in an avian, female heterogametic system

**DOI:** 10.1101/2021.08.17.456618

**Authors:** Ana Catalán, Justin Merondun, Ulrich Knief, Jochen B.W. Wolf

## Abstract

The evolution of genetic sex determination is often accompanied by degradation of one of the proto sex chromosomes. Male heterogametic systems have evolved convergent, epigenetic mechanisms restoring the resulting imbalance in gene dosage between diploid autosomes (AA) and the hemizygous sex chromosome (X). Female heterogametic systems (AA_f_ ZW_f_, AA_m_ ZZ_m_) tend to only show partial dosage compensation (0.5 < Z_f_:AA_f_ < 1) and dosage balance (0.5<Z_f_:ZZ_m_<1). The underlying mechanism remains largely elusive. Here, we quantified gene expression for a total of 15 male and female Eurasian crows (*Corvus (corone) spp.*) raised under common garden conditions. In addition, we characterized aspects of the regulatory landscape quantifying genome-wide ATAC-seq and 5mC methylation profiles. Partial dosage compensation was explained by female upregulation of Z-linked genes accompanied by increased chromatin accessibility on the female Z chromosome. 5mC methylation was strongly reduced in open chromatin-regions and GC islands and showed chromosome-, but no sex-specific variation. With the exception of the pseudo-autosomal region (PAR), female upregulation of gene expression was evenly spread across the Z chromosome without evidence for regional epigenetic regulation, as has for example been suggested for the male hypermethylated region (MHM) in chicken. Our results support the hypothesis that partial dosage compensation in female heterogametic systems is subject to chromosome-wide, epigenetic control mediated by differential chromatin accessibility between the sexes.

## Introduction

The evolution of chromosomal sex-determination appears to be accompanied by degradation of one of the proto sex chromosomes resulting in male (m) or female (f) heterogameity (Bachtrog, 2013; Graves, 2006). In the former case, the set of heteromorphic sex chromosomes resides in males (m: XY; f: XX), in the latter case in females (f: ZW; m: ZZ). In both systems, the heterogametic sex is left with only half of the original copy number of orthologues genes on the shared sex chromosome (X or Z, respectively). In the absence of compensatory mechanisms, this change in gene dosage cuts ancestral transcript levels in half (Gelbart and Kuroda, 2009; Graves, 2016; Schield et al., 2019) and induces an imbalance in gene regulatory networks with genes from diploid autosomes (X_m_:AA_m_ = Z_f_:AA_f_ = 0.5) (Bachtrog, 2013; Charlesworth, 1996; Mank, 2009). In the homogametic sex, the sex-chromosomal to autosomal gene dosage (XX_f_:AA_f_; ZZ_m_:AA_m_) remains, in principle, unaffected (Vicoso and Bachtrog, 2011).

Chromosome-level haploinsufficiency, and even dosage change in single genes, will alter stochiometric ratios of interacting proteins, generally with strong deleterious fitness effects (Birchler et al., 2005; Rice and McLysaght, 2017). A single-copy sex chromosome thus constitutes a serious peril for the hemizygous sex and is expected to prompt an evolutionary response (Charlesworth, 1996; Gu and Walters, 2017; Ohno, 1967). Indeed, male heterogametic systems as divergent as fruit flies and mammals have independently evolved molecular mechanisms that function to restore expression balance (Livernois et al., 2012; Samata and Akhtar, 2018). While the degree of compensation and the underlying mechanisms differ across taxa (Gu and Walters, 2017; Mank, 2009), epigenetic regulation and changes in chromatin structure are common themes (Duncan et al., 2018; Klemm et al., 2019; Park and Kuroda, 2001). For example, in mammals one of the X chromosomes is randomly targeted for inactivation in the homogametic sex by nucleosomal condensation impeding transcription (Graves, 2014). The active X chromosome is then hyper-transcribed to equalize expression levels to the (diploid) autosomal dose (Nguyen and Disteche, 2006). In *Drosophila*, in contrast, hyperactivation is limited to heterogametic males where relaxation of the chromatin structure on the X restores expression levels to the default double-dosage (Samata and Akhtar, 2018). In *Caenorhabditis elegans* a hitherto unknown mechanism upregulates expression levels in both sexes. While this restores dosage in heterogametic males, expression overshoots in females and requires a secondary mechanism to return gene expression on the X to parity with the autosomes (McDonel et al., 2006). X-specificity is achieved by DNA motifs guiding the dosage compensation complex (Ercan et al., 2007; Jans et al., 2009). A common feature of the abovementioned XY systems is that dosage compensation appears to be initiated at specific sites along the X chromosome (Ercan et al., 2007; Drosophila: Gelbart and Kuroda, 2009; mice: Lin et al., 2007).

Global compensatory mechanisms restoring hemizygous to autosomal balance are not ubiquitous. In heterogametic ZW systems, females tend to only show partial compensation, a discrepancy that may follow from basic evolutionary principles (Naurin et al., 2010). This lack of full dosage compensation is observed across a diverse set of female heterogametic taxa including lepidopterans (Catalán et al., 2018; Gu et al., 2017; Zha et al., 2009), snakes (Schield et al., 2019; Vicoso et al., 2013a) and birds (Itoh et al., 2010, 2007) – the focal taxon of this study for which most information is available. The basal avian group of ratites, where sex chromosomes still recombine and degradation has been trapped at an arrested stage, still show Z:A parity across the largely homomorphic sex chromosomes (Adolfsson and Ellegren, 2013; Vicoso et al., 2013b). Other species in the avian tree of life differ in the degree of morphological and genomic divergence of the Z-chromosome and the size of the diploid pseudo-autosomal region (Rutkowska et al., 2012; Zhou et al., 2014). Yet, all species with heteromorphic sex chromosomes investigated to date show only partial compensation for dosage differences.

Following (Gu et al., 2017), we differentiate between dosage balance and dosage compensation. The former refers to parity in expression between male and female genes located on the shared sex chromosome (X_m_:XX_ff_ or Z_f_:ZZ_m_ = 1). The latter refers to expression parity between sex-chromosomes of either sex and the ancestral, non-degenerated proto sex chromosome. In the absence of ancestral information, autosomal levels are generally used as a proxy (X_m_:AA_m_ = XX_f_:AA_ff_ = 1 or Z_f_:AA_f_ = ZZ_m_:AA_m_ = 1). This proxy seems justified under the conditions of equal expression for autosomal genes between the sexes (AA_f_:AA_m_ = 1) and lower intra-autosomal variance relative to variance between autosomes and the sex chromosome. The heterogametic bird species investigated so far tend to show both partial dosage balance (0.5 < Z_f_:ZZ_m_ < 1) and partial dosage compensation (0.5 < Z_f_:AA_f_ < 1). Depending on the species and tissue, f:m expression ratios of Z-linked genes range from 0.67 – 0.81 (Ellegren et al., 2007; Itoh et al., 2010, 2007; Uebbing et al., 2013; Wolf and Bryk, 2011). Z:A expression ratios similarly range from 0.5 – 0.7 in females, but reach equity in homogametic males (Ellegren et al., 2007; Uebbing et al., 2013; Wolf and Bryk, 2011).

This range between 0.5 (pure dosage effect) and 1 (full compensation) requires a mechanism conveying partial compensation and balance. The nature of this mechanism, however, remains elusive. In some species, a specialized region of greater dosage compensation near the male hypermethylated locus (MHM) would suggest localized epigenetic mechanisms on the Z chromosome analogous to XY systems (Arnold et al., 2008; Melamed and Arnold, 2007; Sun et al., 2019). Revision of the MHM in chicken using chromosome-wide DNA-methylation tiling arrays, however, did not confirm a clear relationship between sex-specific methylation and regional dosage balance (Nätt et al., 2014). Moreover, in the majority of investigated species patterns of f:m gene expression appear to be homogenous along the Z-chromosome and do not call for a regional epigenetic mechanism (Itoh et al., 2010; Sun et al., 2019; Wolf and Bryk, 2011). Thus, it has been hypothesized that partial dosage compensation in birds is most likely regulated on a gene-by-gene basis (Mank and Ellegren, 2009), possibly in combination with a chromosome-wide epigenetic mechanism (this study).

The genomic tools to quantify gene expression, assay epigenetic profiles and quantify chromatin accessibility are readily available (Jordan et al., 2019). Yet, comprehensive studies investigating epigenetic mechanisms and chromatin structure in birds are rare (Fishman et al., 2019; Xu and Millar, 2020) and in the context of dosage compensation near-absent (Nätt et al., 2014; Sun et al., 2019). In this study, we quantified gene expression in two somatic tissues, spleen and liver, for a total of 15 male and female Eurasian crows (*Corvus (corone) spp.*) raised under common garden conditions. In addition, we characterized the genome-wide chromatin accessibility profile by means of ATAC-seq and generated methylation data to assay chromosomal 5mC profiles similarly known to affect gene expression (Lee et al., 2020; Lindner et al., 2021). Linking these three data types allowed us to gain insight into possible epigenetic mechanisms underlying partial dosage compensation.

## Results

### Chromosome-wide patterns of gene expression, chromatin accessibility and 5mC methylation

The interrogation of open chromatin states and 5mC methylation levels are expected to covary with a set of genomic features such as chromosome length, gene length and GC content, known to differ widely among avian chromosomes (O’Connor et al., 2019). Therefore, categorical comparisons between the long Z chromosome (75 Mb) and all autosomes including micro-chromosomes (4.88–154.33 Mb; RefSeq GCF_000738735.5) are only meaningful if these factors are statistically controlled for. Accordingly, subsequent analyses focusing on differences between autosomes and the Z chromosome included these features as constitutive covariates where appropriate. Our measures of interest are also expected to vary in different parts of the chromosome (e.g. genic, intergenic). We therefore additionally considered different scales of integration: entire chromosomes (chromosome-wide), actively expressed genes including +/- 20kb of putative regulatory regions (genes) and flanking regulatory regions without the gene body (regulatory). Statistical results for all measures are summarized across these levels in a **Supplementary Model** file. **Supplementary Table A** summarizes all f:m and Z:A ratios (AA_f_:AA_m_, Z_f_:ZZ_m_, Z_f_:AA_f_, ZZ_m_:AA_m_,), both as raw estimates and parameter estimates with confidence intervals controlling for covariates. For chromosome comparisons, the Z chromosome was considered as a whole including the pseudo-autosomal region (PAR). Effects of the PAR were then isolated in a second step.

#### Gene expression

RNA-seq data from liver and spleen confirmed partial dosage compensation and dosage balance, as previously demonstrated for brain tissue in the same species (Wolf and Bryk, 2011). In both liver and spleen tissue, expression levels were statistically undistinguishable between autosomal genes of both sexes and genes located on the male Z chromosome (AA_m_ ≅ ZZ_m_ ≅ AA_f_ ≅ 1) (**Figure AA, Supplementary Model A**). Expression of Z-linked genes in females was significantly reduced relative to males (parameter estimate Z_f_:ZZ_m_= 0.63/0.62 for liver/spleen respectively; same nomenclature hereafter) and relative to genes on female autosomes (Z_f_:Aa_f_ = 0.68/0.68). In both cases, confidence intervals of expression ratios were between 0.5 and 1 confirming the hypothesis of partial dosage compensation through female upregulation of Z-linked genes.

#### Chromatin structure

We hypothesized that upregulation on the female’s Z chromosome might be under epigenetic control. We therefore performed an ATAC-seq experiment which provides information about nucleosome spacing and chromatin accessibility along the genome (Buenrostro et al., 2013). The density of ATAC-seq signals is expected to scale positively with the accessible proportion of DNA. Accordingly, peak density was taken to reflect global accessibility of a given chromosomal region. The height of individual peaks was used as a measure reflecting local accessibility of putative regulatory regions (Chen et al., 2016). We present the results of both measures in turn.

##### Peak density

Analogous to gene expression, chromosome-wide autosomal peak density was statistically undistinguishable between the sexes. On the Z-chromosome, however, peak density was significantly reduced in females relative to males (Z_f_:ZZ_m_= 0.66/0.62). Variation among autosomes was high and raw Z:A ratios were broad, yet overall with significantly lower values in females (Z_f_:AA_f_ = 0.66/0.60, ZZ_m_:AA_m_ = 0.95/1.02, **Supplementary Model B1**, **Supplementary Figure SE)**. To establish the link between gene expression patterns and chromatin accessibility more closely, we next restricted our analysis to peaks located in and around genes. Setting a threshold of 20 kb up- and downstream of the gene model, we scored a median of 6–7 linked peaks per expressed gene **(Supplementary Figure SF).** Gene-centered peak density closely recapitulated the chromosome-wide pattern (**Figure AB**, **Supplementary Model B2, Supplementary Figure G**). Peak density on the single Z copy in females reached about ¾ of the density of autosomes (Z_f_:AA_f_= 0.77/0.73, ZZ_m_:AA_m_= 0.93/0.97) and of the male Z chromosome (Z_f_:ZZ_m_= 0.78/0.61) suggesting a non-linear relationship between the number of accessible chromosome regions and ploidy. The pattern also remained stable when excluding gene bodies considering only putative regulatory regions up- and downstream of the expressed gene (**Supplementary Model B3)**. In summary, peak density closely resembled the partial compensation pattern of gene expression across all scales of integration.

##### Peak height

The average height of ATAC-seq peaks identified across autosomes did not differ between the sexes and was undistinguishable from levels on the male Z chromosome. Peak height in females, however, was significantly elevated on the Z when compared to males (Z_f_:ZZ_m_=1.10/1.16) and when compared to the autosomes (Z_f_:AA_f_=1.12/1.06, **Supplementary Figure SE, Supplementary Model C1**). These differences in peak height were stable when considering only peaks associated with actively expressed genes (**Figure AC, Supplementary Model C2**) and peaks residing in putative regulatory regions up- and downstream of the expressed genes (**Supplementary Model C3, Supplementary Figure G**). This effect was not merely due to few outlier regions with extreme values of peak height, but was consistent across the peak height distribution. In all quantiles, peak height exceeded autosomal values (**Supplementary Figure SH)**. The effect was also stable across varying intervals around genes including 1–15kb up- and downstream of the gene model (**Supplementary Figure SI**). Moreover, higher accessibility in females was also observed at the level of ATAC-seq read coverage excluding possible artifacts of the peak calling algorithm. As expected, f:m coverage ratio was centered around 1 for autosomes (AA_f_:AA_m_ ≅ 1.03/1.03), but was elevated on the female Z beyond the expectation from chromosomal copy number of 0.5 (Z_f_:ZZ_m_ = 0.65/0.65**, Supplementary Model C4**). Overall, this observation suggests that open chromatin is most accessible on the hemizygous, female Z chromosome.

#### 5mC methylation

DNA methylation constitutes another candidate regulatory mechanism underlying partial dosage compensation. To assess its role, we generated whole-genome bisulfite sequencing data of one female and one male individual for both liver and spleen. First, we investigated the relationship between 5mC methylation with gene expression and chromatin accessibility. Analogous to the chromatin features described above, we assayed 5mC methylation at different scales: chromosome-wide, gene-centered (gene body +/- 20 kb flanking region), putatively regulatory regions in CpG islands and overlapping open chromatin regions of genes (ATAC-seq peaks), both including and excluding the gene body (**Supplementary Figure SJ**). As expected, methylation levels were high chromosome-wide (proportion of global methylation: 0.56/0.58) and around genes (0.63/0.63), but much reduced (and bimodal) in CpG islands (0.05/0.06) and in open chromatin regions including (0.02/0.07) or excluding the gene body (0.02/0.03) (**Supplementary Figure SK**). The Z chromosome as a whole exhibited slight hypermethylation (0.60/0.60) relative to macrochromosomes of similar size (chromosomes 1– 21; 0.56/0.57), with minor hypermethylation observed in spleen across microchromosomes (chromosomes 22–28; 0.59/0.62). Important in the context of dosage compensation and dosage balance, hypermethylation on the Z was near-identical between the sexes (**Supplementary Models D1**). When restricting the analyses to genic regions, there was still no difference between the two sexes, but hypermethylation was no longer visible (**Supplementary Models D2).** In CpG islands, the Z chromosome was strongly hypermethylated in liver indicating either an excess of hypermethylated islands on chromosome Z or a reduction of hypermethylated islands on the autosomes **(Supplementary Figure SK)**. Male to female methylation levels remained at parity (**Supplementary Model D3**). In open chromatin regions, there was no difference between the sexes, only among chromosomes. Open-chromatin regions in Z-linked genes appeared to be hypermethylated in liver and hypomethylated in spleen for both sexes **(Supplementary Figure SL, Supplementary Models D4 & D5)**. Overall, these results indicate a tissue- and chromosome-specific 5mC methylation landscape in crows. They provided no evidence for a possible role of 5mC methylation contributing to dosage compensation or dosage balance. Yet, the fact that 5mC methylation levels were correlated with open-chromatin features across sexes and chromosomes (**Supplementary Figure SM**) allows for a possible indirect contribution.

**Figure A.**
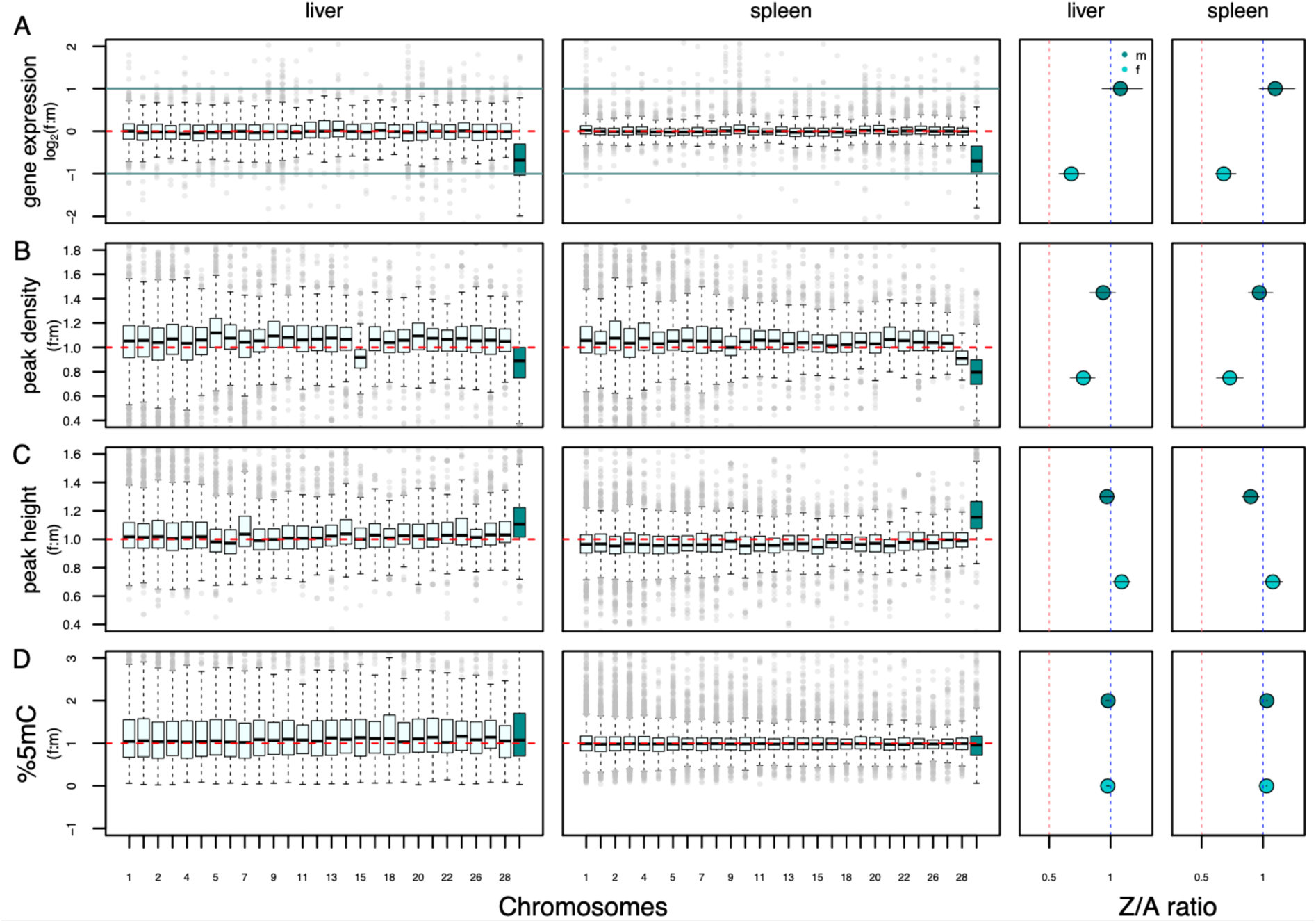
**Gene-centered patterns of dosage effects for gene expression, chromatin accessibility and 5mC methylation.** The two leftmost columns show f:m ratios by chromosome for liver and spleen with the Z chromosome highlighted in color. Horizontal red dashed lines indicate an equal f:m ratio. Boxes encompass interquartile range across genes, whiskers extend to 1.5 x the interquartile range. The two rightmost columns show Z-chromosome to autosomal (Z:A) ratios separately for females (Z_f_:AA_ff_, light shading) and males (ZZ_m_:AA_m_, dark shading). Here, red vertical lines mark a ratio of Z/A=0.5 and blue vertical lines mark a ratio of Z/A = 1. Bars in dotplots represent 95% confidence intervals of parameter estimates of statistical models controlling for co-variates (**Supplementary Models**) drawn from 1,000 bootstraps. **(A)** Log_2_(f:m) gene expression. **(B)** Female to male (f:m) ATAC-seq peak density. **(C)** Female to male (f:m) ATAC-seq peak height. **(D)** Female to male (f:m) methylation percentages.

### Patterns of gene expression, chromatin accessibility and 5mC methylation along the Z chromosome

#### Identification of the pseudo-autosomal region (PAR)

Next, we examined the distribution of dosage effects along the Z chromosome. In a first step, we identified the pseudo-autosomal region of the crow genome for use as a “diploid control”. Combining information from f:m differences in sequencing coverage, heterozygosity levels along the Z chromosome and orthology with other avian species, we defined a PAR region of ∼688 kb from the proximal end of the Z chromosome (**Figure C, Supplementary Figure SN**). In the PAR, we identified 36 annotated genes, 14 of which were expressed in liver and 16 in spleen (for a .gff file see **Supplementary Table SB**). We then calculated f:m ratios inside (PAR) and outside the PAR (nonPAR) for gene expression, peak density and peak height.

Analogous to the chromosome comparisons above, statistical results are summarized in the **Supplementary Model** file for genes and putative regulatory regions. Note that due to the low sample sizes of genes on the PAR, standard errors were large and power was low. **Supplementary Table C** summarizes all PAR:nonPAR ratios between females and males both as raw estimates and parameter estimates with confidence intervals controlling for covariates.

**Figure B.**
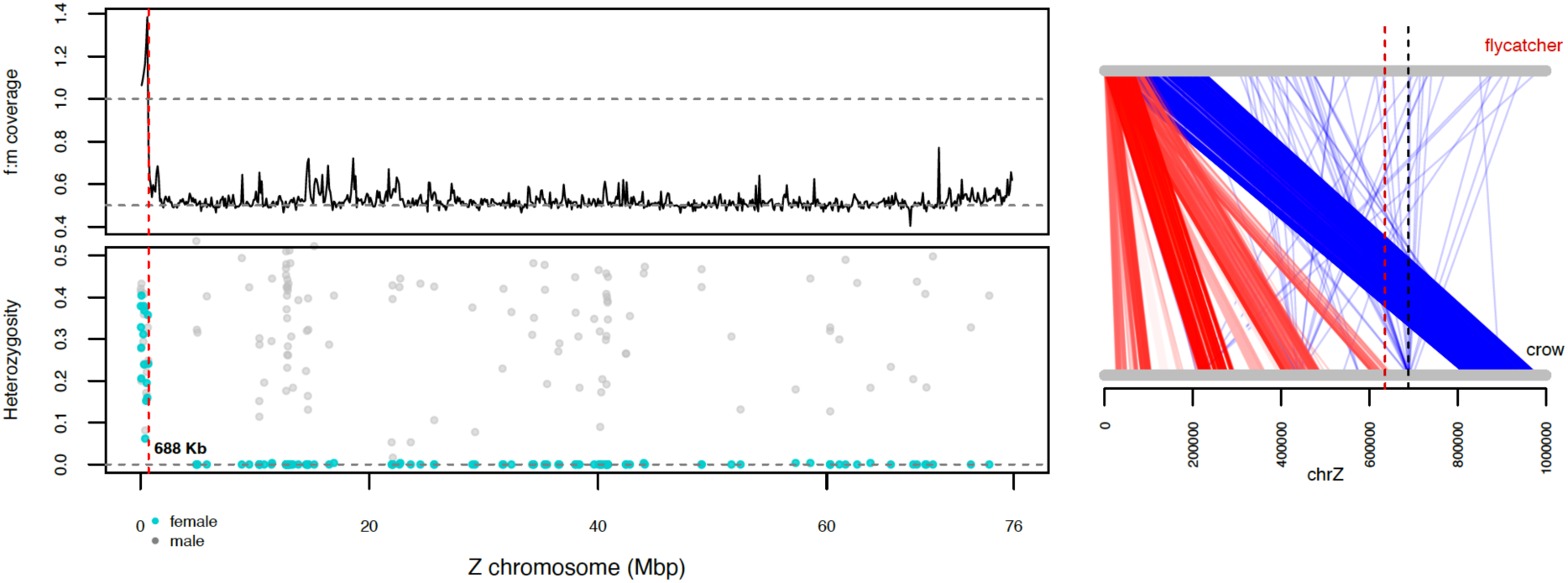
Identification of the pseudo-autosomal region (PAR) in the European crow. Left upper panel: f:m genome coverage along the Z chromosome. Left bottom panel: heterozygosity levels across the Z chromosome in females (light blue) and males (gray). A drop in female coverage and heterozygosity at 688 kb defines the end of the PAR. Right panel: 1 Mb of Z chromosome alignment between the flycatcher and the European crow. Lines between the Z represent orthologous regions found between the two species. Red: protein coding genes. Blue: non-coding regions. The vertical dashed line marks the end of the PAR region in crows. Vertical dashed red line marks the PAR as identified in the flycatcher (Smeds et al., 2014).

#### Gene expression

As expected by gene dose, all genes expressed in the PAR had equal expression levels in females and males in both tissues (PAR_f_:PAR_m_=1.01/1.01, **Supplementary Figure SO**). Genes located in the remaining part of the Z chromosome (nonPAR) showed the expected pattern of reduced expression in females (nonPAR_f_:PAR_f_=0.55/0.78, **Supplementary Model E1**). For the non-PAR part of the Z chromosome we further categorized genes into fully compensated (-0.25 < Z_f_:ZZ_m_ < 1), partially compensated (-0.25 > Z_f_:ZZ_m_ > -0.75) and uncompensated (-0.75 > Z_f_:ZZ_m_ > -1). The number of genes falling into these categories was similar in liver and spleen. For each tissue, 147/159 genes (28.1/29.2%) were fully compensated, 99/122 genes (19.9/18.3%) were partially compensated and the majority of 277/350 genes (52.9/52.5 %) showed no evidence of compensation according to this definition. The number of genes expressed in both tissues was not enriched for compensation status (Fisher’s exact test *P*-value: 0.43) indicating no preference for compensation of housekeeping genes.

Subsequent sliding window analysis along the Z chromosome with a window size of 688 kb (PAR regions equivalent) or 1 Mb and a step size of 344 kb or 100 kb, respectively, revealed no clustering of compensated genes (**Figure CA**). Not a single window was enriched for compensated genes (Fisher’s exact tests with multiple testing correction, **Supplementary Table SD)**. The same result was obtained when the enrichment analysis was done using windows containing 15 genes (mimicking the number of expressed genes on the PAR; **Supplementary Table SD**). A lack of regional concentration of compensated genes was further corroborated by a lack of autocorrelation in Z_f_:ZZ_m_ values (**Supplementary Figure SP**).

#### Peak density

Under the hypothesis that gene expression patterns are governed by chromatin accessibility, we did not expect regionally clustered variation in ATAC-seq peak density and height on the Z chromosome either. As for gene expression, peak density in the PAR was comparable between sexes and autosomal values of the nonPAR region in males. In females, peak density in the nonPAR was lower than in the PAR across all levels of integration (**Supplementary Models E2-E3**). A sliding window analysis (window size of 688 kb and 344 kb shift) identified no region on the Z enriched for a higher density of peaks in females. In males, three non-overlapping regions on the Z were identified with higher peak density in both organs (dark blue areas in **Figure CC**). In spleen, ten additional regions were identified (**Figure CB, Supplementary Tables SE)**. Overall, these results are consistent with homogeneously reduced peak density outside of the PAR region in females (**Figure CB**).

#### Peak height

The height of peaks associated with genes also did not differ between sexes in the PAR region and was close to a ratio of one (**Supplementary Model E4, Supplementary Figure SO**. For the remaining part of the Z chromosome, peak height was significantly elevated in females relative to the syntenic nonPAR region in males. This pattern was stable across tissues and across genes or regulatory regions alone (**Supplementary Models E4-E5**). Of all 1761/1770 peaks shared between females and males, 90/78 peaks were female-biased and 16/17 were male-biased in liver and spleen, respectively, although after multiple testing significance was lost (**Supplementary Table SE)**. Using a sliding window approach (window size 688 kb, shift 344 kb), we uncovered two regions with significantly higher peaks for females in both liver and spleen (light blue ares in **Figure CC**) and an additional 15 non-overlapping regions present in spleen only (**Supplementary Table SF**), suggesting tissue specific regulation of chromatin accessibility. Overall, these analyses confirm increased peak height in the single copy region of the Z chromosome in females relative to males without much evidence of local clustering (**Figure CC**).

#### 5mC methylation – no evidence for a male hypermethylated region

In chicken, two male hypermethylated regions (MHM) have been identified on the Z chromosome, in which genes appear dosage compensated as a result of gene expression arrest through male methylation (Sun et al., 2019; Wright et al., 2015). In the crow data, we found no significant regional enrichment of male methylation. At base-pair resolution, we observed several localized regions of DNA methylation that differed between the sexes along chromosome Z, including the PAR. We identified high autocorrelation within these regions among many of the 5mC positions for both sexes, suggesting sexually differentiated methylated blocks interspersed along the Z chromosome **(Supplementary Figure SQ)**. Yet, there was no consistent pattern of exclusively male nor female-biased blocks. We then replicated the above analyses for methylation of genes and CpG islands (see feature dataset **Supplementary Figure SR**). Regions of sexual differentiation differed compared to the analysis at base pair resolution, but likewise identified no male (nor female) hypermethylated region.

**Figure C.**
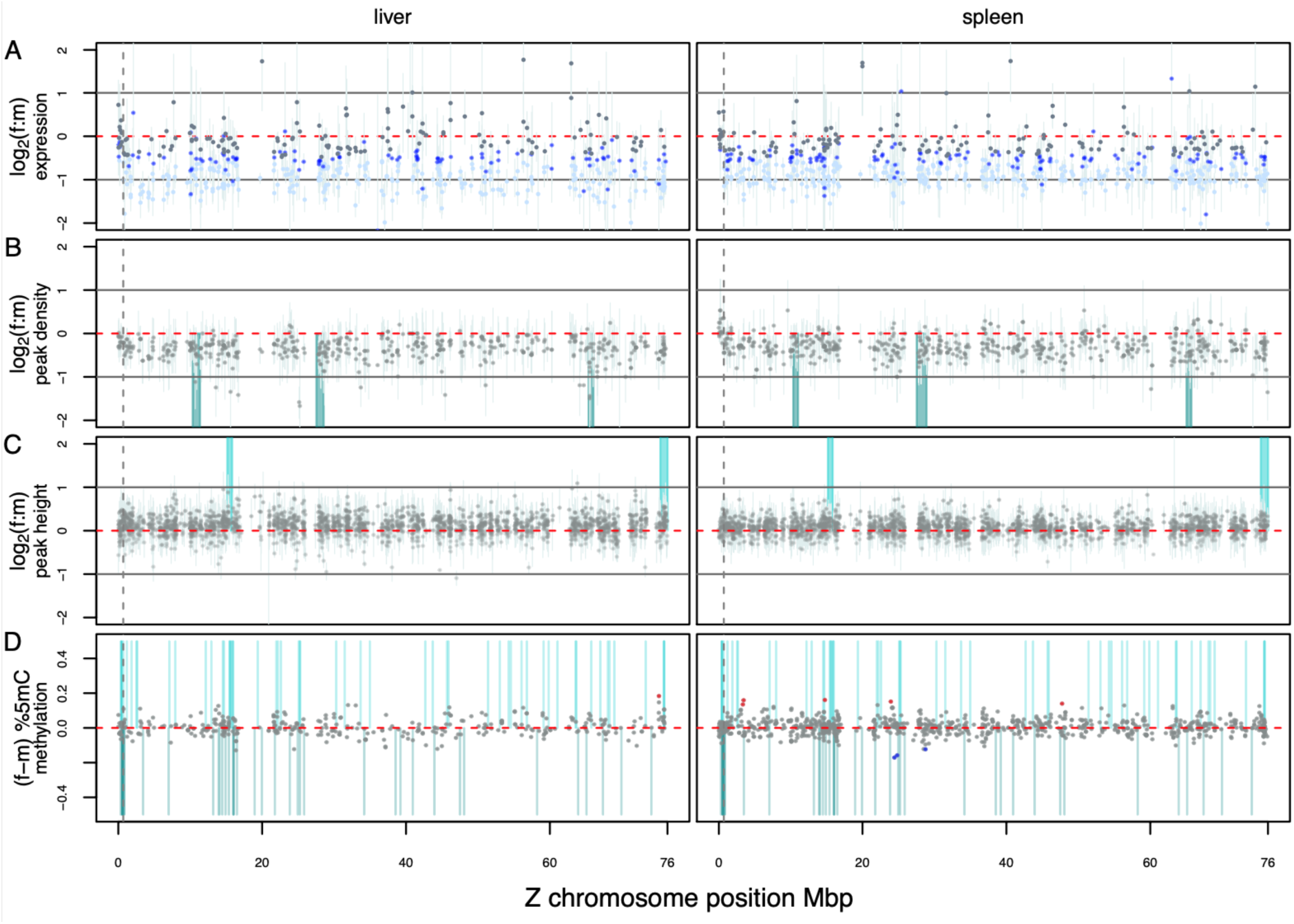
Mean log_2_(f:m) along the Z for (**A**) gene expression (**B**), peak density (**C**), peak height (**D**) and methylation. **A–C**: For each estimate, 95% confidence intervals were calculated as the median of bootstrapped female and male values (10,000 times) of gene-centered estimates. Shaded vertical areas represent regions shared between organs with significant enrichment of compensated genes in males (dark blue, values < 0) or females (light blue, values > 0). **A:** Gray dots represent dosage compensated genes, royal blue genes with partial compensation and light blue genes with no compensation.

## Discussion

In this study, we quantified chromosome and sex-specific variation in gene expression (RNA-seq), and used two methods to interrogate the underlying epigenetic variation including chromatin features (ATAC-seq peak density and height) and levels of 5mC methylation (WGBS). We interpret our findings in light of dosage compensation and dosage balance.

### Gene expression: Z_f_ < (ZZ_m_ = AA_f=m_)

Gene expression data provided evidence for incomplete dosage compensation (Z_f_:AA_f_ < 1) and dosage balance (Z_f_:ZZ_m_ < 1) on a background of autosomal parity between the sexes (AA_f_ = AA_m_). The expression pattern of Z_f_ < (ZZ_m_ = AA_f=m_) has been observed in many ZW chromosome systems such as birds, snakes and *Heliconius* butterflies, irrespective of the age of the non-degraded sex-chromosome (Catalán et al., 2018; Gu et al., 2017). Deviations from this pattern have been suggested for moths (Smith et al., 2014) and livebearer fish (Darolti et al., 2019). Our statistical analyses controlling for confounding genomic features show that gene expression on the single-copy part of the female Z chromosome was significantly increased over the expectation of gene dose alone. This was observed relative to the expression of genes residing on the Z in females and males (0.5 < Z_f_:ZZ_m_ < 1), relative to overall expression of autosomal genes when compared to the Z (0.5 < Z_f_:AA_f_ < 1), and relative to genes on the dual-copy, pseudo-autosomal region of the female Z chromosome (0.5 < nonPAR_f_:PAR_f_ <1). Using autosomal levels of gene expression as an approximation for ancestral, diploid expression levels, this result argues for upregulation of genes located on the hemizygous Z chromosome in females during the course of sex chromosome evolution. This conclusion is consistent with previous point estimates from other avian species exceeding Z_f_:AA_f_ expression ratios of 0.5 for the majority of genes (Ellegren et al., 2007; Itoh et al., 2010; Julien et al., 2012; Uebbing et al., 2013; Wolf and Bryk, 2011), but see (Chen et al., 2020).

### Epigenetic dosage effects: from pattern to mechanism?

Epigenetic variation is an important determinant of variation in gene expression (Ando et al., 2019). Sex- and chromosome-specific regulation of epigenetic variation thus constitutes a plausible mechanism to equalize differences in gene dose (Duncan et al., 2018; Graves, 2014; Park and Kuroda, 2001; Samata and Akhtar, 2018). We here quantified broad-scale patterns of chromatin structure and 5mC methylation serving as proxies for epigenetic mechanisms with direct and indirect effects on transcription (Klemm et al., 2019; Lindner et al., 2021). This approach is based on the rationale that these two epigenetic methods are, in essence, functionally related to gene expression. Several lines of evidence support this hypothesis. ATAC-seq peak height is known to translate into higher chromatin accessibility (Corces et al., 2018), and other studies have seen a correlation between the number of chromatin accessible sites and the level of gene expression (Marlétaz et al., 2018; Werner et al., 2018). A regulatory role of open chromatin is not necessarily restricted to the close vicinity of genes, as regulatory elements can be found megabases away from the target gene (Hug et al., 2017). Local variation in 5mC methylation is of undoubted importance for gene expression (Veselovska et al., 2015), and has previously been discussed in the context of dosage compensation (Richard et al., 2017). Also, for several of the chromatin accessibility metrics and methylation levels considered here, we observe an expected relationship with broad-scale patterns of gene expression. Clearly, in the absence of a detailed regulatory map in crows inferring process from (broad-scale) pattern is a leap of faith, and results need to be interpreted with caution. Many of the open chromatin and methylated regions will not bear any direct functional relationship to gene expression as has been shown by other studies (Klemm et al., 2019). Still, we believe that the overall patterns described in the study do contain meaningful information. To mitigate potential biases and reduce potential background noise concealing functionally active genomic regions, we quantified patterns along a gradient of increased functional resolution ranging from the coarse category of chromosomes, to genes and fine-scaled overlap of ATAC-seq peaks with 5mC methylation data.

An additional challenge of interpreting patterns of epigenetic variation across sexes and chromosomes is the influence of confounding variables. Unsurprisingly, variation in both the ATAC-seq and 5mC methylation data was in part explained by chromosomal features such as chromosome length, gene length or GC content. All of these variables are closely related to recombination rate and are known to influence many aspects of genome biology in birds (Nam et al., 2010; O’Connor et al., 2019). For example, in the neighborhood of genes, nucleosome spacing (peak density) and accessibility of nucleosome free regions (peak height) were both influenced by chromosome length and gene length. In short, open chromatin was sparser, but more accessible the longer the genes and chromosomes. The degree of 5mC methylation also depended on chromosome length and GC content with hypomethylated genes residing on longer chromosomes. Clearly, these confounding variables need to be statistically accounted for to obtain sex-specific parameter estimates isolating dose effects between the Z chromosome and the highly variable autosomes. The use of flexible, mixed-effects linear models serves this purpose and has been advocated in the context of dosage compensation before, though has rarely been put into practice (Gu and Walters, 2017). Discrepancies between our model-based estimates and Z:A ratios derived from raw data in this study underline the importance of this aspect (**Supplementary Table A, C**).

### Density of open chromatin regions: Z_f_ < (ZZ_m_ = AA_f=m_)

Patterns of open chromatin density closely mirrored patterns of gene expression. Chromatin accessible regions were spaced at equal rate across male and female autosomes and the male Z chromosome. Density of open chromatin peaks was only reduced on the female Z. Strikingly, density was not reduced to half, but to a ratio in between 0.5 and 1, mimicking the pattern of partial dosage compensation for gene expression. This pattern was stable across scales of resolution: genome-wide, gene-centered and regulatory. This may partly be explained by the fact that the majority of all the identified peaks (55%) resided near expressed genes (+/- 20kb) diminishing the influence of intergenic regions. Nonetheless, near-identical f:m and Z:A ratios and confidence intervals across scales of integration speak for a rather homogenous, chromosome-wide effect. Under the justified assumption that homologous chromosomes share a substantial part of open chromatin regions, this observation does not necessarily require an independent epigenetic mechanism of the paternally inherited, single-copy Z chromosome of females. Z_f_:ZZ_m_ ratios between 0.5 and 1 may simply reflect the proportion of open chromatin peaks constitutively shared between homologous chromosomes. Differentiating between this scenario and active depletion of nucleosome density on the female Z would require quantification of ATAC-seq peaks for each of the male homologues separately.

### Accessibility of open chromatin regions: Z_f_ > (ZZ_m_ = AA_f=m_)

In contrast to gene expression and peak density, peak height was elevated on the female Z chromosome relative to males and autosomes of both sexes. Again, estimates in f:m and Z:A ratios were near-identical across scales of integration pointing towards a chromosome-wide mechanism. We therefore hypothesize that a majority of genes achieve partial dosage compensation and balance through a sex-specific, chromosome-wide mechanism increasing chromatin accessibility throughout the female Z chromosome. A similar chromatin Z-specific remodeling mechanism leading to a permissive chromatin landscape has been suggested in other, non-avian female heterogametic systems (Picard et al., 2019). In principle, such a mechanism is compatible with a gene-by-gene mechanism of compensation, as has previously been suggested for birds (Mank, 2009; Mank and Ellegren, 2009; Warnefors et al., 2017). Gene-to-gene variation in the degree of compensation can still be achieved by variation in the distribution of open chromatin regions and other epigenetic marks. It is, however, incompatible with different levels of compensation strength corresponding to evolutionary strata on the Z chromosome (Wright et al., 2015; Xu et al., 2019b).

### Epigenetic patterns along the Z

Our hypothesis of a chromosome-wide epigenetic mechanism was supported by our analysis of epigenetic patterns along the Z chromosome. To address the question of heterogeneity along the Z chromosome, we performed window-based analyses. First, we isolated the homogametic PAR region, which proved to be comparable in size and gene content to other passerines (Xu et al., 2019a; Zhou et al., 2014). Parameter estimates of expression levels, peak density and peak height on the female PAR were statistically comparable to genes on the male PAR. Outside of the female PAR, we identified a reduction of gene expression and peak density, but an increase in peak height. Due to the small size of the PAR, confidence intervals were large, but estimates reflected the overall patterns observed between the hemizygous Z chromosome and the autosomes. Approximately 47% of the expressed genes that were classified to be compensated were randomly distributed along the Z. For peak density, the sliding window analysis uncovered three shared regions in liver and spleen that showed significantly lower peak density when compared to the PAR in females. These regions could contain male specific regulation centers supporting higher transcription of genes beneficial to males. For peak height, two regions on the female Z were significantly elevated compared to the PAR in both tissues, pointing towards highly localized regions of female-bias, which could function as additional coordination centers for gene regulation. Overall, however, these findings are consistent with chromosome-wide regulation of open chromatin and provide no evidence for a localized ‘organization center’ of dosage compensation.

### 5mC methylation: no global nor regional dosage effect

A lack of regional regulation of Z-linked gene expression was likewise evident when considering 5mC methylation levels. As expected, methylation differed strongly among levels of integration. Gene bodies were most strongly methylated, followed by CpG islands and regions of accessible chromatin. Chromatin accessibility is expected to show a strong negative relationship with DNA methylation (Taudt et al., 2016), as DNA methylation hinders the anchoring of the transcription machinery and tightens up the chromatin structure (Duncan et al., 2018; Moore et al., 2013). Despite these expected genome-wide correlations, patterns of DNA methylation levels were neither sex-specific, nor did they show any clear relationship to the pattern of partial dosage compensation observed for gene expression and ATAC-seq data. Whole-genome methylation data rather portrayed a conserved methylomic landscape on the Z chromosome between sexes. This is consistent with previous reports in chicken and the white-throated sparrow (Sun et al., 2019). Yet, we found no evidence for male-specific hypermethylation on the Z chromosome, as suggested for chicken (Arnold et al., 2008; Melamed and Arnold, 2007; Sun et al., 2019) and other species in the Galloanserae clade (Wright et al., 2015). Instead, we identified regions of increased differences in methylation consisting of hypo- and hypermethylated sites in both sexes. The lack of a MHM region in the European crow further supports a growing body of evidence that male-specific hypermethylated regions are far from ubiquitous across Aves (Itoh et al., 2010; Uebbing et al., 2013), and these regions are also absent in whole-genome bisulfite data in the white-throated sparrow (Sun et al., 2019). The lack of a MHM region further indicates that in the European crow, the DNA methylation landscape on the Z chromosome does not contribute to regional sex-specific gene expression to achieve dosage compensation. This is consistent with mechanisms detected in zebra finch, where methylation is autonomous from dosage compensating mechanisms in brain tissue (Diddens et al., 2021). Instead, we hypothesize that the female’s Z has evolved a chromatin environment that increases permissiveness along the entire chromosome and that gene-specific control of expression is fine-tuned by specific chromatin accessible regions.

### Conclusions

This study provides evidence for partial dosage compensation and dosage balance in crows as a result of female upregulation of Z-linked gene expression. It further sheds light on the potential underlying epigenetic mechanisms suggesting a chromosome-wide effect of increased chromatin accessibility on the hemizygous Z chromosome. 5mC methylation appeared to play no role, nor did regional ‘centers’ of regulation on the Z chromosome, such as the male hypermethylated region, find any support. The data also provided no hint at an influence of discrete genomic regions differing in the age of recombination cessation (evolutionary strata (Xu et al., 2019a). Dedicated, comparative dosage compensation studies across the avian tree will be needed to see whether signals of gene expression and epigenetic patterns still reflect the ghost of past evolutionary strata. In summary, this work highlights the role of chromatin dynamics in the process of dosage compensation and constitutes a first step towards a functional understanding of partial dosage compensation mechanism in birds.

## Material and methods

### Taxonomic considerations

The two taxa considered here – carrion crows (*Corvus (corone) corone*) and hooded crows (*Corvus (corone) cornix*) – have been considered as separate species by some (Linnaeus, 1758; Parkin et al., 2003). Recent genome-wide evidence, however, rather supports treatment as a single species with two segregating colour morphs (Knief et al., 2019; Knijff, 2014; Poelstra et al., 2014; Vijay et al., 2016). Relevant for the context of this study, previous work has also found near-identical gene expression profiles (Warmuth et al., 2021) with marked differences only for genes expressed in melanocytes underlying plumage divergence (Poelstra et al., 2015; Wu et al., 2019). For the purpose of this study, we therefore consider carrion and hooded crows as members of the same species.

### Data set and sample collection

In May 2014, crow hatchlings of an approximate age of 21 days were obtained directly from the nest (Weissensteiner et al., 2015). Hooded crows (*C.* (*corone*) *cornix*) were sampled in the area around Uppsala, Sweden (59°52’N, 17°38’E), and carrion crows (*C.* (*corone*) *corone*) in the area around Konstanz, Germany (47°45N’, 9°10’E). A single individual was selected from each nest to avoid any confounding effects of relatedness. After transfer of carrion crows to Sweden by airplane, all crows were hand-raised indoors at Tovetorp field station, Sweden (58°56’55”N, 17°8’49”E). When starting to feed by themselves, they were released to large roofed outdoor enclosures (6.5 x 4.8 x 3.5 m), specifically constructed for the purpose. All crows were maintained under common garden conditions in groups of a maximum of six individuals separated by sub-species and sex. In October 2016, at an age of approximately 2.5 years, individuals were euthanized by cervical dislocation. Tissues were dissected from eight females (3 *corone*, 5 *cornix*) and seven males (4 *corone*, 3 *cornix*). Tissue for RNA extraction was conserved in RNAlater (ThermoFisherScientific) and stored at -80°C. Tissue designated for ATAC-seq was flash frozen at -80°C immediately after dissection. Metadata on all individuals can be retrieved in **Supplementary Table SJ**.

*Regierungspräsidium Freiburg* granted permission for the sampling of wild carrion crows in Germany (Aktenzeichen 55-8852.15/05). Import into Sweden was registered with the *Veterinäramt Konstanz* (Bescheinigungsnummer INTRA.DE.2014.0047502) and its Swedish counterpart *Jordbruksverket* (Diarienummer 6.6.18-3037/14). Sampling permission in Sweden was granted by *Naturvårdsverket* (Dnr: NV-03432-14) and *Jordbruksverket* (Diarienummer 27-14). Animal husbandry and experimentation was authorized by *Jordbruksverket* (Diarienummer 5.2.18-3065/13, Diarienummer 27-14) and ethically approved under the Directive 2010/63/EU on the Protection of Animals used for Scientific Purposes by the *European Research Council* (ERCStG-336536).

### RNA-seq

#### Data generation and processing

RNA-seq reads were generated for liver and spleen. Frozen tissue was disrupted with a TissueRuptor (Qiagen, Hilden, Germany), and total RNA was extracted with the RNeasy Plus Universal Kit (Qiagen, Hilden, Germany). RNA quantity and quality were determined with an Agilent Bioanalyzer 2100 (Agilent Technologies, Santa Clara, CA). RNA-seq libraries were prepared from 500 ng total RNA using the TruSeq stranded mRNA library preparation kit (Illumina Inc, Cat# 20020594/5) including polyA selection. The library preparation was performed according to the manufacturers’ protocol (#1000000040498). Fifty base pair single-end reads were generated on a HiSeq2500, v4 sequencer (Illumina). Sequencing was performed by the SNP&SEQ Technology Platform in Uppsala, Sweden.

Base pair quality was assessed with FastQC v0.11.5 (Andrews, 2010) and bases with a Phred quality score < 20 were removed. All reads with a minimum length of 20 bp were kept for mapping. Reads were mapped to the crow genome version 5.6 (NCBI: ASM73873v4) with STAR (Dobin et al., 2013). An annotation lift-over was done from crow genome version 2.5 (Poelstra et al., 2014) to the version 5.6. We used StringTie (Pertea et al., 2015) to predict novel isoforms, that were merged into the annotation lift-over file if not overlapping with already annotated features. Raw read counts per annotated transcript were calculated with HTSeq (v0.9.1) (Anders et al., 2015). Fragments Per Kilobase per Million mapped reads (FPKM) were calculated and normalized with Cufflinks (Trapnell et al., 2013). For empirical summary statistics of central tendency, we filtered out genes if the mean FPKM of the focal gene across all females and across all males was <1. Thus, genes with FPKM > 1 were considered expressed. For all model-based inference, “actively expressed genes” were identified by calculating zFPKM-values for each sample and tissue using the zFPKM R package (v1.10.0) and we considered genes with zFPKM > -3 as expressed (Hart et al., 2013).

#### Quantification of dosage compensation and dosage balance

For each tissue, f:m expression ratios were calculated per chromosome (A1A1_f_:A1A1_m_, A2A2_f_:A2A2_m_, … Z_f_:ZZ_m_) as log_2_(median(Gene_i_FPKM_female_)/median(Gene_i_FPKM_male_)) for each gene. Sex-specific Z:A ratios (Z_f_:AA_f_, ZZ_m_:AA_m_) were calculated as median(log_2_(FPKM of all Z-linked genes)) / median(log_2_(FPKM of all A_i_-linked genes)). The degree of dosage compensation was categorized by partitioning log_2_(f:m) expression of each gene into three categories: (1) compensated: log_2_(f:m) > -0.25 and log_2_(f:m) < 0.25, (2) partially compensated: -0.25 > log_2_(f:m) > -0.5 and 0.25 < log_2_(f:m) < 0.5 and (3) uncompensated: log_2_(f:m) < -0.75 and log_2_(f:m) > 0.75. All log_2_(f:m) > 2 were classified as female or male biased genes.

### ATAC-seq

#### Data generation and processing

We followed an ATAC-seq protocol specifically developed for frozen tissue (Corces et al., 2017). In brief, ∼ 10 mg of cryopreserved liver and spleen tissue samples were ground using a TissueRuptor (Qiagen, Hilden, Germany) in liquid nitrogen and subsequently homogenized in HB Buffer (see buffer recipes in Corces et al. 2017). Cells were counted with a hemocytometer and 50,000 cells were used for transposition. Transposition was carried out for 60 min in a 50µl reaction with 2.5 µl (100nM final) of Tn5 transposase (Illumina Nextera) per sample. Libraries were amplified using the primers published in (Buenrostro et al. 2013). The number of necessary library PCR amplification cycles was determined by qRT-PCR using SYBR Green 10x in DMSO (Sigma-Aldrich) and NEBNext Master Mix 2x (New England BioLabs). DNA concentration of the resulting library was determined with a Qubit 2.0 Fluorometer (Life Technologies, Grand Island, NY) and its quality was assessed using an Agilent Bioanalyzer 2100 (Agilent Technologies, Santa Clara, CA). Libraries that passed the quality controls (bioanalyzer fragment distribution following nucleosomal patterning) were selected for sequencing. Two technical replicates were prepared per sample and sequenced (50 bp paired-end) on an Illumina HiSeq1500 sequencer in an inhouse sequencing facility at LMU Munich and on an Illumina NovaSeq S1 flowcell by the SNP&SEQ Technology Platform in Uppsala.

Sequencing data quality was assessed with FastQC (v0.11.5), and base pairs with a sequencing Phred quality score < 20 were removed. Nextera adapters were trimmed, and reads with a minimum length of 36 bp were selected for further mapping using Cutadapt (v1.9.1) (Martin, 2011). Selected reads were mapped to the crow genome version 5.6. (NCBI accession number: GCA_000738735.4) using Bowtie2 (v2.3.2) (Langmead et al., 2009), with the following parameters: bowtie2 –very-sensitive -X 2000, allowing for fragments of up to 2kb to align. Only properly paired and uniquely mapped reads with a mapping quality score > Q20 were filtered out using samtools (v1.4) (Li et al., 2009). PCR duplicate reads were removed with Picard tools (v2.0.1).

#### Quality control

Following guidelines of ATAC-seq quality control assessment, we included the following indicators to judge data quality: (1) *Insert size distribution* following a nucleosomal spacing periodicity **(Supplementary Figure SA)** (2) *Spearman correlation coefficients* between technical replicates **(Supplementary Figure SB)** (3) The *number of usable reads* (4) The *fraction of reads in peaks* (FRiP) as calculated with deepTools -plotEnrichment **(Supplementary Figure SC)** and (5) *ATAC-seq coverage enrichment at transcription start sites* (TSS), **(Supplementary Figure SD)**. More details on quality control are provided in the **Supplement**. Technical replicates that survived the quality check were merged by biological replicate for all subsequent analyses. A summary of the usable reads can be found in **Supplementary Table SG.**

#### ATAC-seq peak calling

Differences in read coverage among samples can influence the sensitivity and specificity of peak identification. To avoid calling higher peak density in samples with more read depth, we down-sampled all the libraries after mapping and filtering to the read number of the smallest library. This resulted in ∼60 million workable reads in liver and ∼90 million in spleen for all samples **(Supplementary Table SH)**. Even though we had a third more reads for spleen, we did not detect more peaks in this tissue, corroborating the fact that we reached a saturation peak-detection point. Down-sampling was done with samtools view -s and coverage equality across libraries was checked by calculating the coverage for windows of 10 bp using deepTools - bamCoverage -bs 10 -p max (v2.27.1) (Ramírez et al., 2016). After library normalization, peaks were called using MACS2 (v2.1.0) macs callpeak -t -n -g 1e9 -f BAMPE (Zhang et al., 2008).

#### ATAC-seq analysis

We quantified two metrics to describe chromatin accessibility for each individual: ATAC-seq peak height and peak density. Peak height was defined as the fold-change value estimated with MACS2 (Zhang et al., 2008). Peak density was defined as the number of ATAC-seq peaks present in a given genomic interval divided by the length of the interval. Chromosome-wide peak density was calculated as the number of peaks per chromosome divided by its length. Gene-centered peak density was accordingly normalized to the length of the focal expressed gene. Empirical chromosome-wide estimates of female to male ratios (f:m) for peak height and density were calculated as the median(peak height or density)_female_/median(peak height or density)_male_. Z-chromosome to autosomes ratios (Z:A) for each sex were calculated as median(Z chromosome peak height or density)/median(autosomes peak height or density). For peak-centered peak heights f:m were calculated as f:m = median(Gene_i_ peak height_female_)/ median(Gene_i_ peak height_male_). For gene-centered peak density estimates, f:m ratios were calculated as f:m = Gene_i_ peak density_female_/ Gene_i_ peak density_male_. Gene-centered Z:A ratios were calculated as described for chromosome-wide.

### 5mC methylation

#### Data generation and processing

We generated whole-genome bisulfite sequencing (WGBS) libraries for one female and one male individual for both liver and spleen tissue. Post-bisulfite converted libraries were generated using the Illumina TruSeq Methylation Kit (EGMK91324) following the manufacturer’s protocol at the SNP&SEQ Technology Platform in Uppsala, Sweden. Raw reads were trimmed using Trim_Galore! v0.6.6 (Krueger, 2012), clipping the first 9 bases from the 5’ ends, and the last base of the 3’ end, following standard post-bisulfite converted WGBS library protocols and as indicated by methylation bias plots **(Supplementary Figure SS)**. Base and tile quality was assessed with FastQC v0.11.9 after trimming (Andrews, 2010). Cleaned reads were then aligned to the reference genome version 5.6 (NCBI: ASM73873v4 with default settings, followed by read de-duplication and extraction of CpG methylation state using Bismark v0.23.0 (Krueger and Andrews, 2011). We retained only CpGs on assigned chromosomes (1 – 28 and Z) and removed any positions overlapping transition mutations (C - T, or G - A SNPs), identified from population resequencing data generated from a previous study (discussed below; (Poelstra et al., 2014). We then further subset and excluded any sites overlapping repeats identified with RepeatMasker v4.1.1 (Smit et al., 2013), using the chicken repeat library and not including short simple repeats (-nolow). We then summarized coverage by sample **(Supplementary Figure SS)**. We also created a CpG island track for assembly version 5.6 because of our interest in regulatory regions, using makeCGI (Wu et al., 2010), requiring stringent thresholds of a length of at least 250bp, a GC content > 60%, and an observed to expected CG ratio > 75%, resulting in 18,526 islands (**Supplementary Table SI**).

#### Population resequencing reanalysis

We reanalyzed population resequencing data for 15 male hooded (*C. (corone) cornix*) and 15 male carrion (*C. (corone) corone*) crows from an existing study (Poelstra et al., 2014) to identify transition SNPs which could confound WGBS methylation calls (*e.g.* a TG genotype at a CG site will incorrectly be called as non-methylated, it is therefore best practices to exclude C – T and A – G sites (Barrow and Byun, 2014). In short, resequencing data was trimmed with BBtools v38.18 (Bushnell, 2021) and aligned to the crow reference genome using BWA v0.7.17 (Li and Durbin, 2009). Duplicates were removed with GATK v4.1.4.1 (Auwera et al., 2013) and variants were called with Freebayes v1.3.2 (Garrison and Marth, 2012) without population priors. SNPs were filtered with bcftools (Li, 2011), excluding INDELs and sites exhibiting the strongest allelic balance (AB), depth (DP), strand bias (SRP, SAP) and read placement (EPP) biases, identified from the top and bottom 2.5% of sites for each metric (only the bottom 2.5% were removed for SRP, SAP, and EPP because only low scores indicate bias). We furthermore removed sites with a quality (QUAL) below 20, a genotype quality (GQ) below 20, and required at least 75% of individuals to share a site with at least a genotype depth of 3 reads. We then retained only C – T or A – G SNP sites using bash.

#### Genome-wide data analyses

We calculated the proportion of methylated to non-methylated sites considering individual 5mC calls only if covered by at least 5 reads. Methylation data was further only considered if a feature of interest (see below) contained at least three 5mC calls, and was scorable for both male and female tissue-specific libraries. DNA methylation analysis was divided into five distinct feature-based analyses to determine the relative impacts of putatively increasing regulatory importance **(Supplementary Figure SJ)**. First, we considered chromosome-wide levels, with the entirety of each chromosome as a feature. Second, we analyzed positions within genes, including each gene and its 20 kb up- and downstream sequence as a feature. We then examined levels within CpG islands, with each island as a distinct feature. Our fourth and fifth analyses analyzed DNA methylation within open chromatin that was shared between males and females (requiring peaks to overlap by at least 75% between sexes to be considered). As peaks differed between liver and spleen, a separate coordinate file was created for each tissue, which was then overlapped with the tissue-specific DNA methylation data. Specifically, the fourth analysis analyzed DNA methylation in open chromatin that either overlapped or was within 20 kb upstream or downstream of a gene. The fifth analysis only included the open chromatin upstream and downstream of a gene, not the peaks overlapping the gene itself.

#### Identification of a male hypermethylated region

Two male hypermethylated (MHM) regions have been identified in Galloanserae on the Z chromosome, although only one region has been found in other avian clades (Sun et al., 2019). Here, we explored the possibility of an MHM region in the European crow by examining base-pair resolution DNA methylation divergence and genomic autocorrelation between sexes and across tissues. We started our search by blasting the known MHM sequences (Sun et al., 2019) against the hooded crow reference, but detected no hits on the Z chromosome. We hypothesized that if the European crow harbored an MHM, it would exhibit male-specific hypermethylation and strong genomic autocorrelation within that region. To test this, we used the same filtered 5mC calls as above, but instead kept each individual position as a feature, and retained only sites shared between both sexes. We then visualized divergence (both f-m and log2(f/m)) along the Z using ggplot2 (Wickham, 2016). To detect regions of high autocorrelation in each sex, we sampled 50 consecutive methylation positions and calculated Spearman correlations of 5mC levels between the first and second half of sites. We then proceeded to examine the remainder of the Z chromosome in tiled 50 base-pair windows. We classified blocks of differentiation as regions exhibiting methylation differences greater than 25% between the sexes and Spearman correlations greater than 0.3, merging any regions within 10 kb. We repeated this process except with a larger 250 base-pair block, and arrived at the same results.

#### Identification of the pseudo autosomal region (PAR)

We took three approaches to identify the PAR in the European crow. (1) Genomic regions on the Z belonging to the PAR are expected to have f:m_coverage_ of approximately 1 for whole-genome shotgun sequencing data, genomic regions outside the PAR of approximately 0.5. To estimate sex-specific coverage along the Z chromosome we used 150bp paired-end Illumina sequencing data from one female and one male crow with NCBI bioproject_id: PRJNA192205 (Poelstra et al., 2014). Reads were mapped with BWA mem (v0.7.17) (Li and Durbin, 2009) to version 5.6 of the crow genome (NCBI: ASM73873v4). Only concordant paired reads with mapping quality >20 were kept using samtools view -u -h -q 20 -f 0×2 (Li et al., 2009). Read duplicates were removed with Picard tools and repeats were masked and filtered using RepeatMasker and the chicken library (v4.0.7) (Smit et al., 2015). Coverage was calculated with bamcoverage deepTools (v2.5.1) (Ramírez et al., 2016) in 50kb windows for the Z chromosome and chromosome 1 and 2 which are similar in size and GC content: bamCoverage --binSize=50000 --normalizeUsingRPKM -p 2. Differences in chromosomal coverage were assessed as follows: the coverage value of each 50 kb window along chromosomes 1,2 and Z was divided by the median coverage of these autosomes. This was done separately for the female and male sample. The normalized coverage was used to calculate log_2_(f:m_coverage_). (2) In addition to the coverage-based approach, we assessed synteny with the PAR of collared flycatcher (*Ficedula albicollis*; (Smeds et al., 2014). We aligned the Z of the flycatcher genome version 1.5 (Kawakami et al., 2014) using the genome aligner lastZ (v1.04.00) (Harris, 2007) and selected only those contigs with > 90% identity. Anchors to homologous regions between the species were plotted in R (R Core Team, 2015) (3) In heterogametic females, we expect to find heterozygous sites in the PAR as opposed to regions outside the PAR, where no heterozygosity should be found. We used GoldenGate genotype data from the Z chromosome (N = 129 SNPs) collected on 522 hooded and carrion crows and their hybrids (data described in Knief et al. (2019)). For each SNP, we calculated the observed heterozygosity in females (N = 277 individuals) and males (N = 245 individuals). We transferred SNP positions to crow genome version 5.6 and observed 15 heterozygous SNPs in females on the Z chromosome between 27,730 and 687,785 bp.

### Statistical analysis

#### Linear models

We fitted linear mixed-effects models with a Gaussian error structure using the lme4 (v1.1-27.1) and lmerTest (3.1-3) R packages (Bates et al., 2015; Kuznetsova et al., 2017). We transformed the data to match normality (dependent variables: FPKM and ATAC peak height were log_2_-transformed, ATAC peak number was square-root-transformed and percent 5mC was arcsine-square-root-transformed). It has been shown that Gaussian models are robust and that violating the normality assumption has only minor effects on parameter estimates and P-values (Knief and Forstmeier, 2021). For each dependent variable, we fitted whether the gene/ATAC peak/methylation window was located on an autosome or the Z chromosome (factor with two levels) and the sex of the individual (factor with two levels) as interacting fixed effects. We controlled for gene length, chromosome length and GC content where appropriate and Z-transformed these covariates. Individual ID and gene ID nested in chromosome ID were modeled as random effects. For the methylation data set, we had only one female and one male sequenced. Because of that, we did not fit individual ID as a random effect. Results should therefore be interpreted with caution, as we are unable to separate sex from individual effects. Exact model specifications are provided in **Supplementary Models**. We used the effects (v4.2-0) R package (Fox, John, Sanford Weisberg, 2011) to obtain the four parameter estimates (autosomes in females, autosomes in males, Z in females, Z in males). We calculated ratios on these parameter estimates and bootstrapped 95% confidence intervals using the bootMer() function in lme4 with 1,000 replicates. These ratios are reported in **Supplementary Table A**, model estimates can be found in **Supplementary Models.** Fixed effect estimates were considered significant when P < 0.05.

For chromosome-wide peak density, we additionally fitted mixed-effects models with a quasipoisson or negative binomial error structure (log-link), using peak number per chromosome as our dependent variables and the same fixed and random effects as above, except that we included chromosome-length as an offset term (Zuur et al., 2009). For this, we used the glmmTMB (v1.1.2) (Brooks et al., 2017) package in R. Exact model specifications are provided in **Supplementary Models** and ratios calculated on parameter estimates are reported in **Supplementary Table A**.

In a second set of models, we restricted the data to the Z chromosome and separated the PAR from the rest of the Z (nonPAR), such that we estimated four parameters (nonPAR in females, nonPAR in males, PAR in females, PAR in males). We controlled for gene length (Z-transformed) by fitting it as a covariate and included individual ID and gene ID as random effects. We used the effects (v4.2-0) R package (Fox & Weisberg 2019) to obtain the four parameter estimates, calculated ratios on these parameter estimates and bootstrapped 95% confidence intervals using the bootMer() function in lme4 with 1,000 replicates. These ratios are reported in **Supplementary Table C**, model estimates can be found in **Supplementary Models.**

#### Identification of female to male ratios along the Z-chromosome

To identify putative clusters of dosage compensated genes on the Z chromosome and to identify regions on the Z enriched or depleted for peak height and peak density, we performed a sliding window analysis. We used a window size of 688 kb (the size of the PAR) and 1 Mb with a sliding step of 344 kb (half PAR size) and 100 kb respectively. Gene expression and ATAC-seq peak metrics on the PAR were used as reference to identify deviating regions along the Z chromosome. A Fisher’s exact test was performed for each window to detect enrichment or depletion of dosage compensated genes along the Z (categories as defined above), with a Bonferroni correction for multiple testing. To detect regions on the Z enriched or depleted or enriched in number of peaks, we additionally used a Wilcoxon test as alternative = “less” and alternative=“greater”. To further survey the Z for potential clustered centers of regulation, we conducted an autocorrelation analysis using the Acf() function from the R package forecast v5.8 (Rob J. Hyndman and Yeasmin Khandakar, 2019). In order to identify peaks with significantly different heights between females and males on the Z, we first identified consensus peaks present in either of the sexes. We selected peaks present in 80% of both female and male samples. Peaks overlapping by more than 50% between individuals were merged into a single peak to avoid peak redundancy across samples. For these consensus-peak coordinates we used extracted peak heights as peak_fold_change computed in MACS2. Only peaks were considered, where at least three individuals had non-zero values. Differences in the peak height distribution between females and males for a given window was tested in R using a Wilcoxon test.

## Acknowledgements

We express our gratitude to Sven Jakobsson for providing the infrastructure for animal husbandry at Tovetorp research station. We would also like to thank Christen Bossu, Jelmer Poelstra and Matthias Weissensteiner for their contribution in obtaining samples. Kristaps Solokovskis, Thomas Giegold, Nils Andbjer, Tamara Volkmer, Barbara Martinschitsch and Luisa Sontheimer provided invaluable support in raising and maintaining the captive crow population. Martin Wikelski, Inge Müller and additional staff from the Max-Planck-Institute for Ornithology in Radolfzell facilitated sampling in Germany and transport to Sweden. We are further grateful to Gabriele Kumpfmüller with whom the ATAC-seq protocol was established. The Swedish sequencing facility is part of the National Genomics Infrastructure (NGI) Sweden and Science for Life Laboratory. We further thank Stefan Krebs at the Gene Center Munich for discussions and sequencing the ATAC-seq libraries.

## Supplementary Text

### Pre-processing of ATAC-seq data

Quality control of ATAC-seq data was performed as described in the main document. Here, we provide details for each of the components. For each library we first examined the insert size distribution which should show a downward laddering pattern reflecting the amount and length of DNA fragments from nucleosome-free region and from regions with associated nucleosomes (Buenrostro et al., 2013). For all libraries, we observed an enrichment of fragments corresponding to the nucleosome-free region that ranged in length from 36 – 130 bp. The second enriched fragment observed, ranged from 150 - 250 bp in both organs, which corresponds to the mononucleosome-bound region (**Figure SA**). The detection of a mononucleosome ensures the boundaries for a successful detection of nucleosome-free regions, where putatively, the regulatory regions lie.

Next, we quantified the correlation between our two technical and biological replicates (8 females, 7 males for liver and spleen). For the technical replicates, Spearman correlation coefficients drawn from ATAC-seq coverage estimates ranged from 0.86 – 0.97 for liver and 0.72 – 0.98 for spleen (**Figure SB**). For biological replicates, Spearman correlation coefficients ranged from 0.87 to 0.95, where correlation was higher within the same sex for both tissues. The number of total usable reads (see methods), varied among samples (**Supplementary Table SB**). Following the ENCODE guidelines for ATAC-seq quality, 50 million fragments are recommended for paired-end experiments. Due to a high correlation between technical replicates, we pooled these for downstream analysis, resulting in ∼60 million reads for the smallest biological sample in liver and ∼92 million reads in spleen.

We also revised the FRiP (Fragments of Reads in Peaks) score values for each pooled sample, which ranged from ∼12 – 51% (**Figure SC**) and which evaluates the number of reads that were mapped into peaks. According the ENCODE guidelines, FRiP values exceeding 0.2 are acceptable. From our sample assessment, only one sample fell below this threshold which was accordingly removed from all subsequent analyses.

We further checked for an enrichment of mapped reads in putative transcription start sites (TSS) of the expressed genes which we defined as genes with FPKM expression levels exceeding 1. The putative TSS was identified as 2kb upstream and downstream of the first nucleotide of the expressed gene. Wig files were produced for each sample and heatmaps showing coverage enrichment around the TSS were done with Deeptools v.3.5.0 (deepTools - computeMatrix -b 2000 -a 2000) (Ramírez et al., 2016). **Figure SD**, shows two examples, one of liver and one of spleen, of coverage enrichment around the TSS (+/- 2kb) (**Figure SD**).

## Supplementary figures

**Figure SA.**
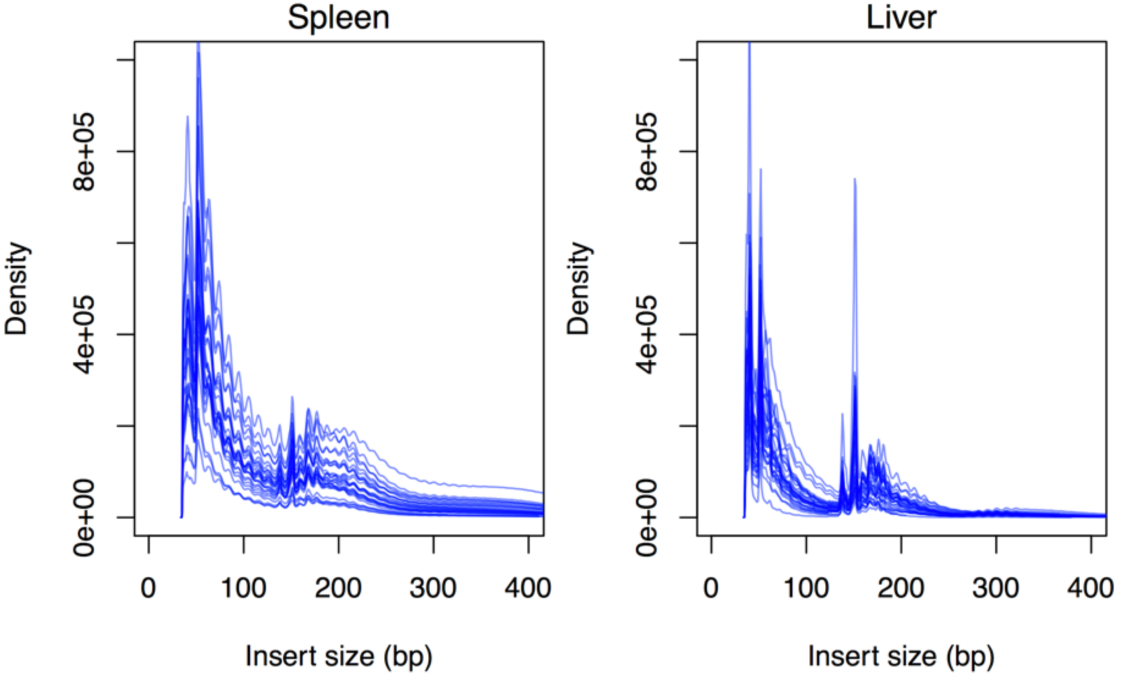
ATAC-seq insert size distribution for all biological replicates for spleen (left) and liver (right). The position of the first nucleosome can be observed between ∼150-250 bp.

**Figure SB.**
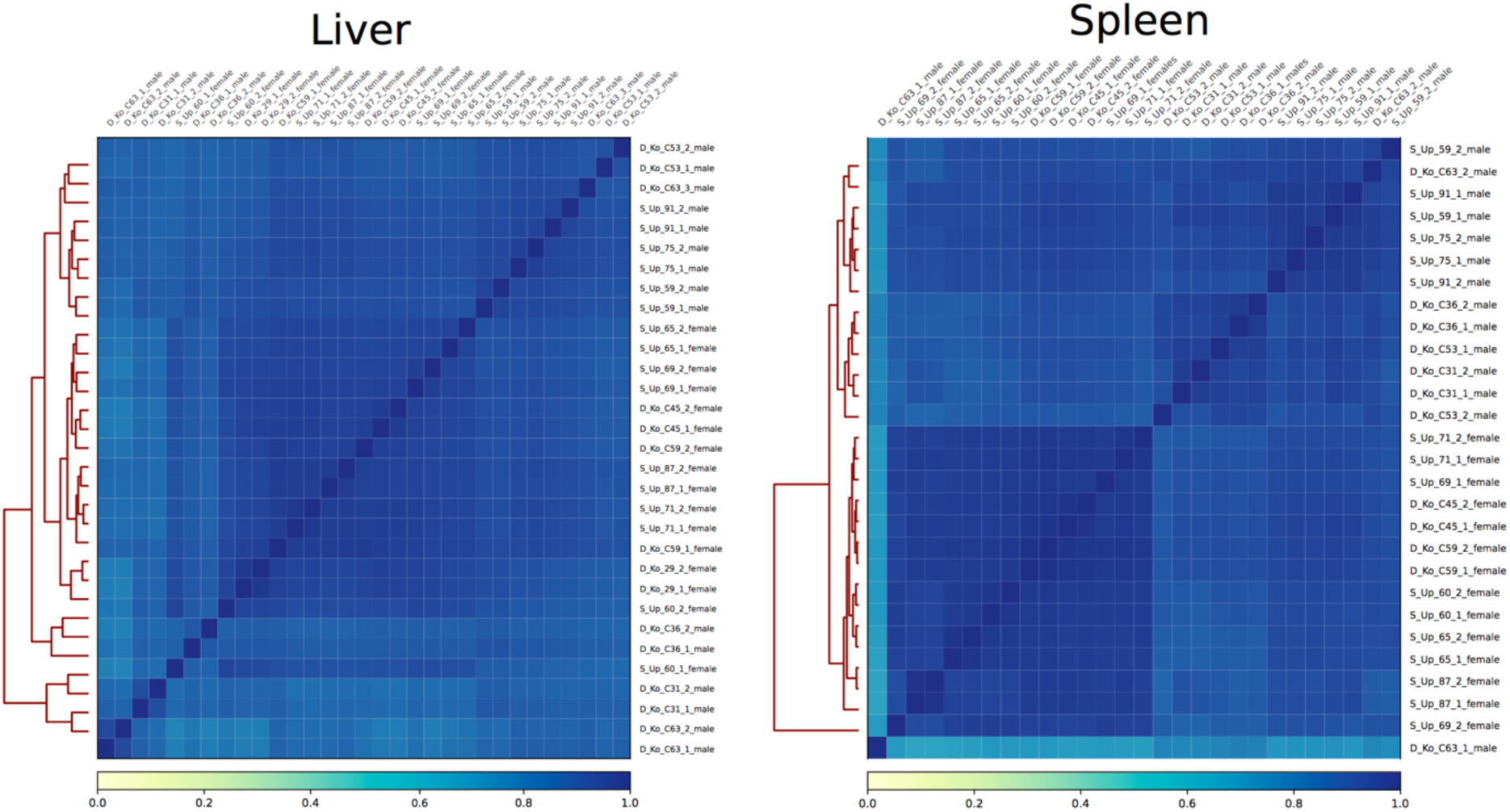
Across sample correlation between technical and biological replicates using genome coverage as the correlation parameter in liver and spleen.

**Figure SC.**
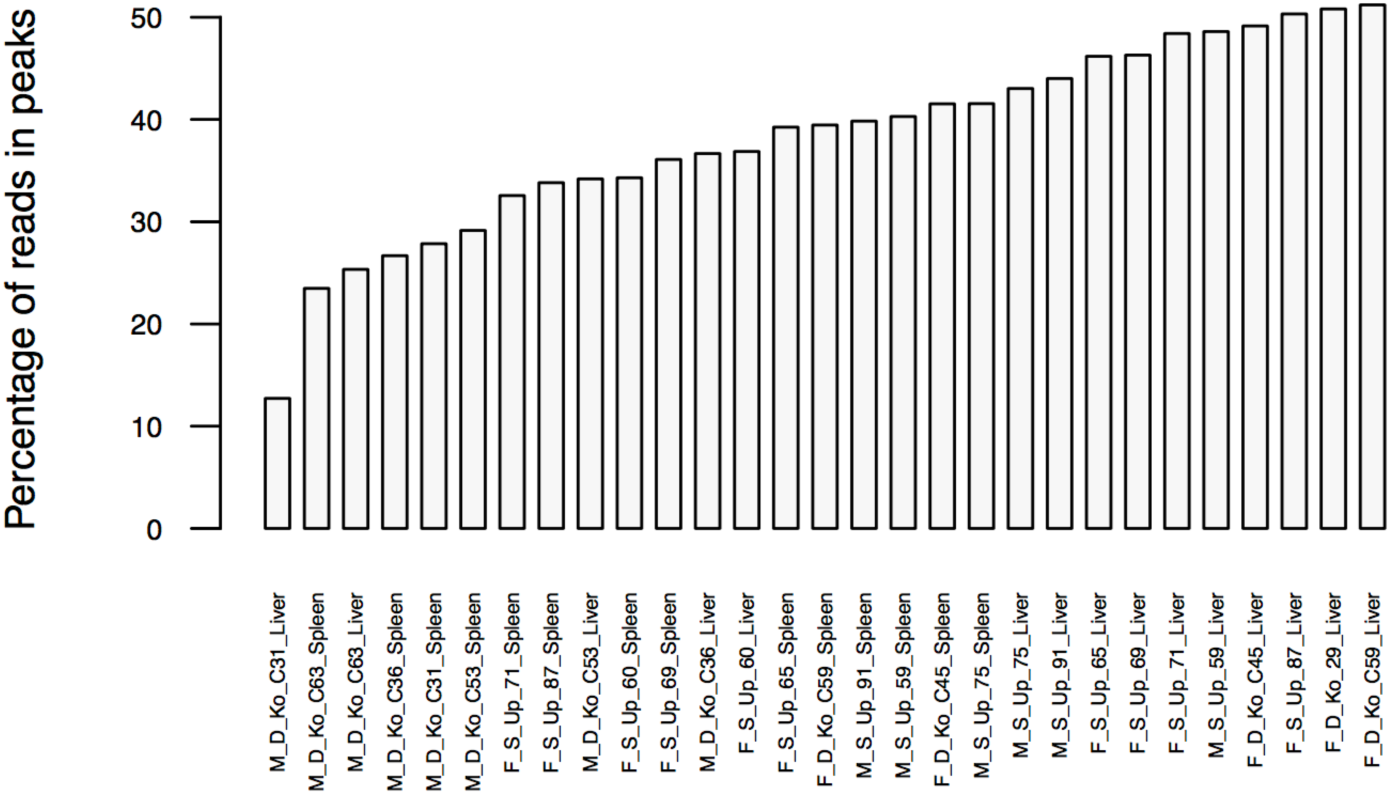
Percentage of reads mapped in reads for pooled samples in liver and spleen. (blue). Vertical dashed lines show the median expression for both density distributions.

**Figure SD.**
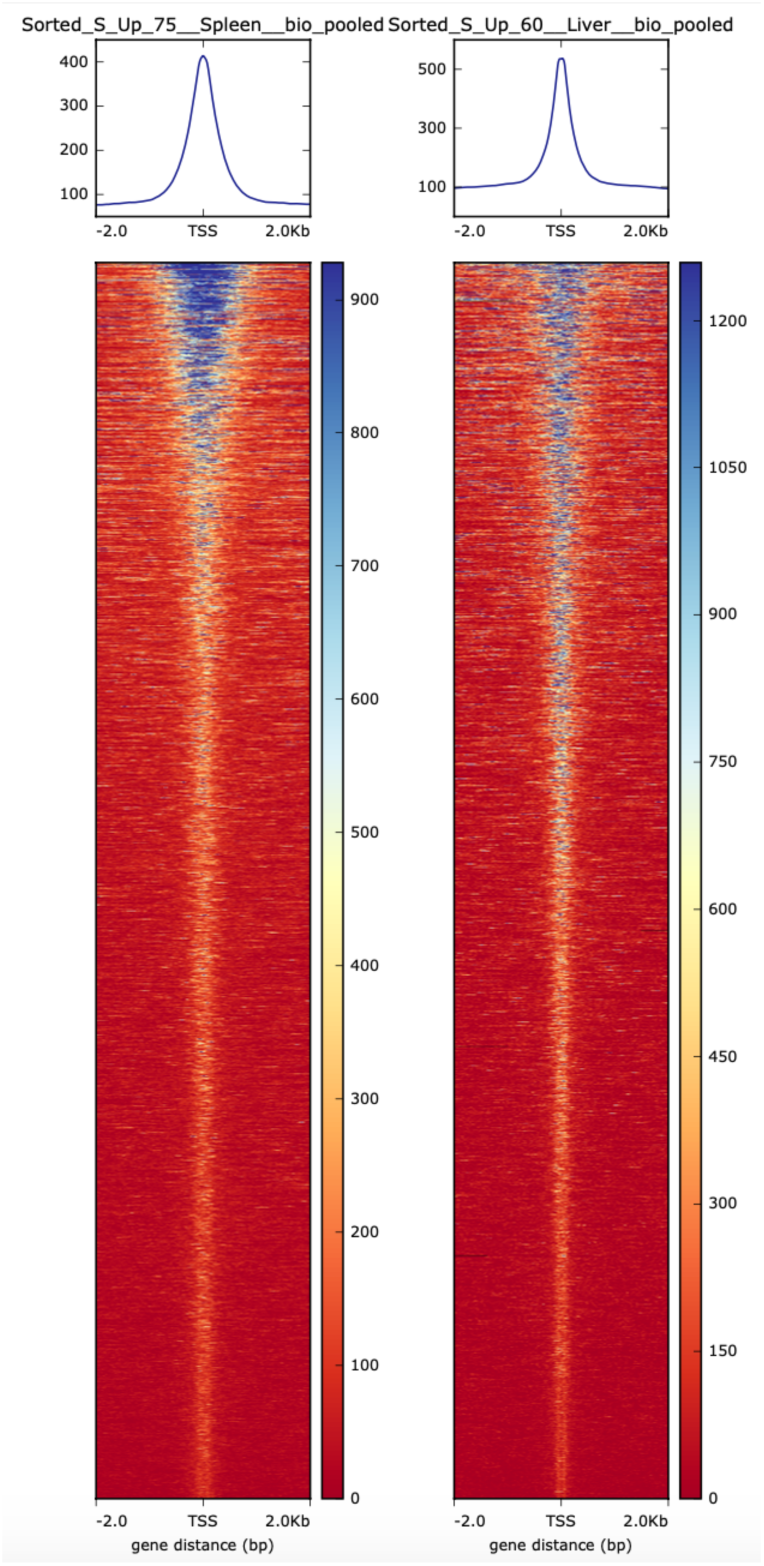
Upper panels: summary of coverage density +/- 2kb of the TSS, in spleen (left) and liver (right). Bottom panels: Enrichment of mapped reads presented as heatmaps along the defined TSS, in spleen (left) and liver (liver).

**Figure SE.**
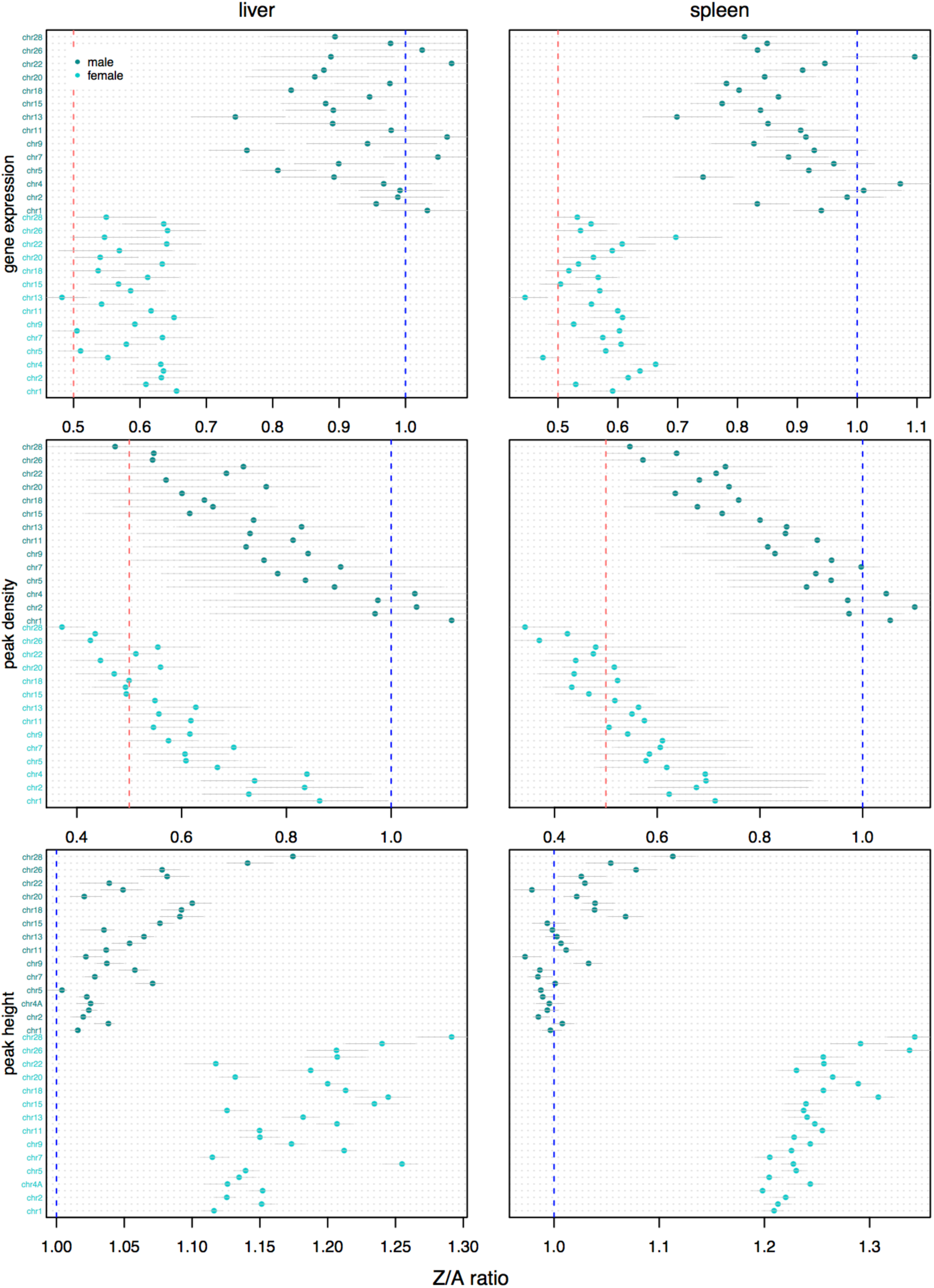
Raw Z:A ratio for gene expression, peak density (chromosome-wide) and peak height (chromosome-wide) shown for each autosome in females (light blue) and males (dark blue). 95% confidence intervals drawn from 10,000 bootstraps.

**Figure SF.**
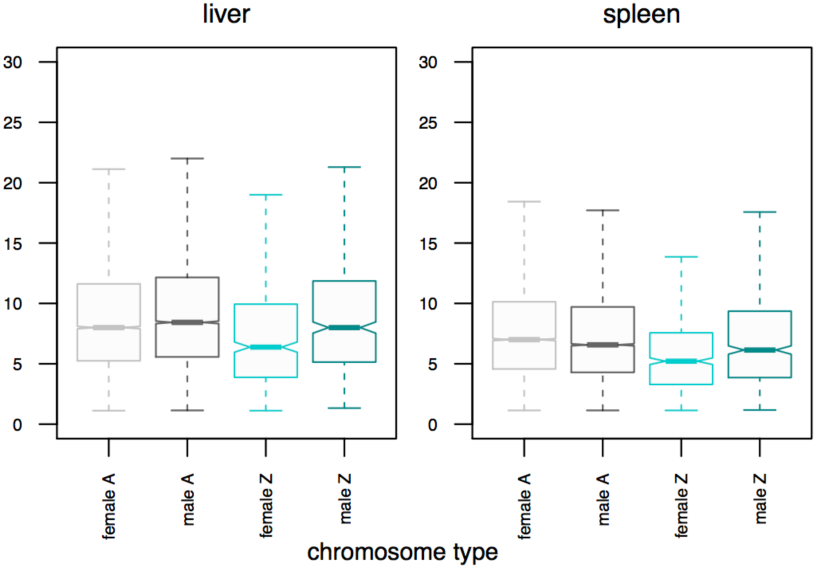
Boxplot showing the distribution of the number of associated peaks per expressed gene in the autosomes and the Z chromosome in a region of 20 kb upstream and downstream of the focal gene. Y-lab indicates the number of peaks associated to a gene.

**Figure SG.**
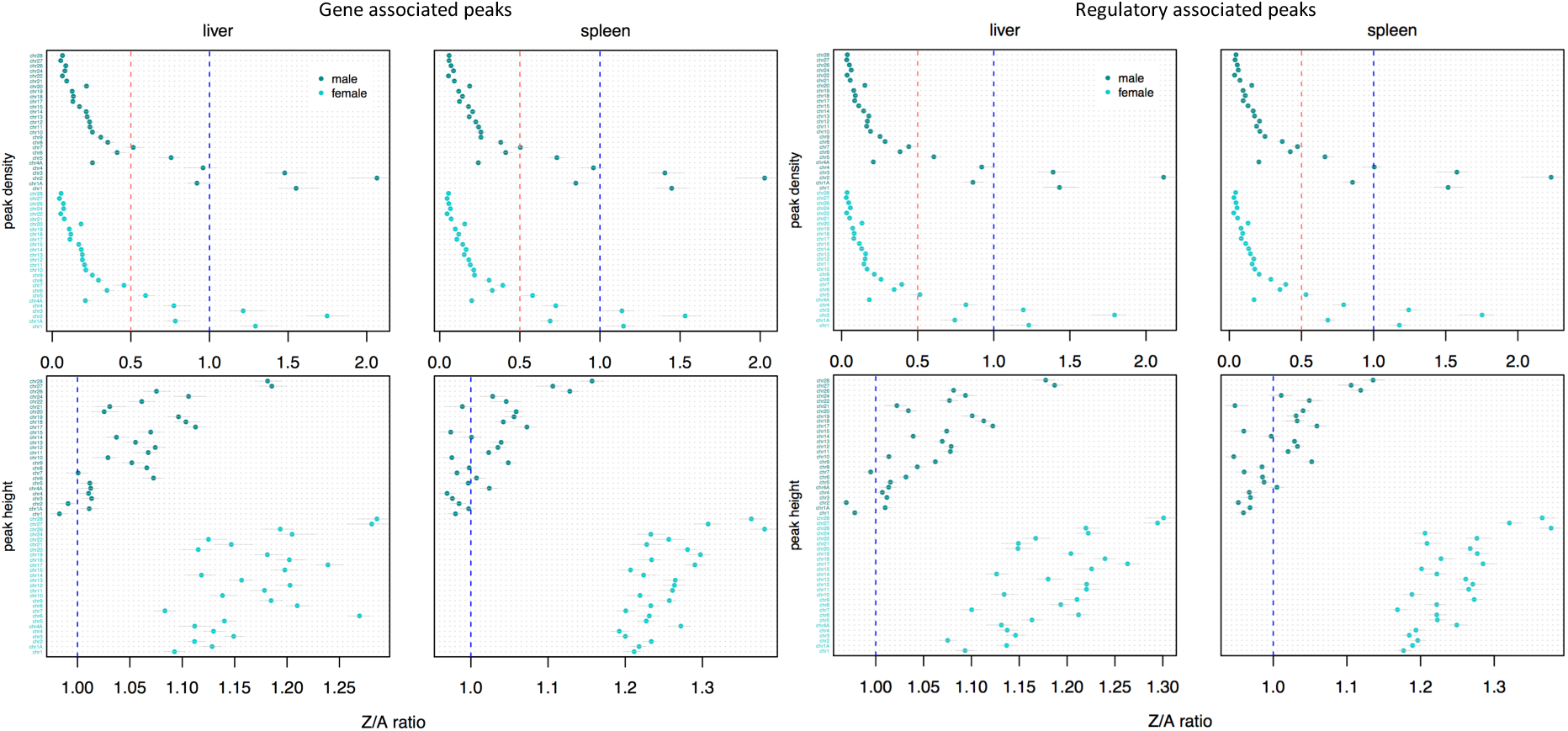
Z:A ratio for peak density and peak height shown for each autosome in females (light blue) and males (dark blue). 95% confidence intervals drawn from 10,000 bootstraps. Left panel: shows gene-centered Z:A ratios. Right panel: shows Z:A only in up- and down- stream regions of expressed genes excluding the gene body. Note the strong dependence on chromosome length.

**Figure SH.**
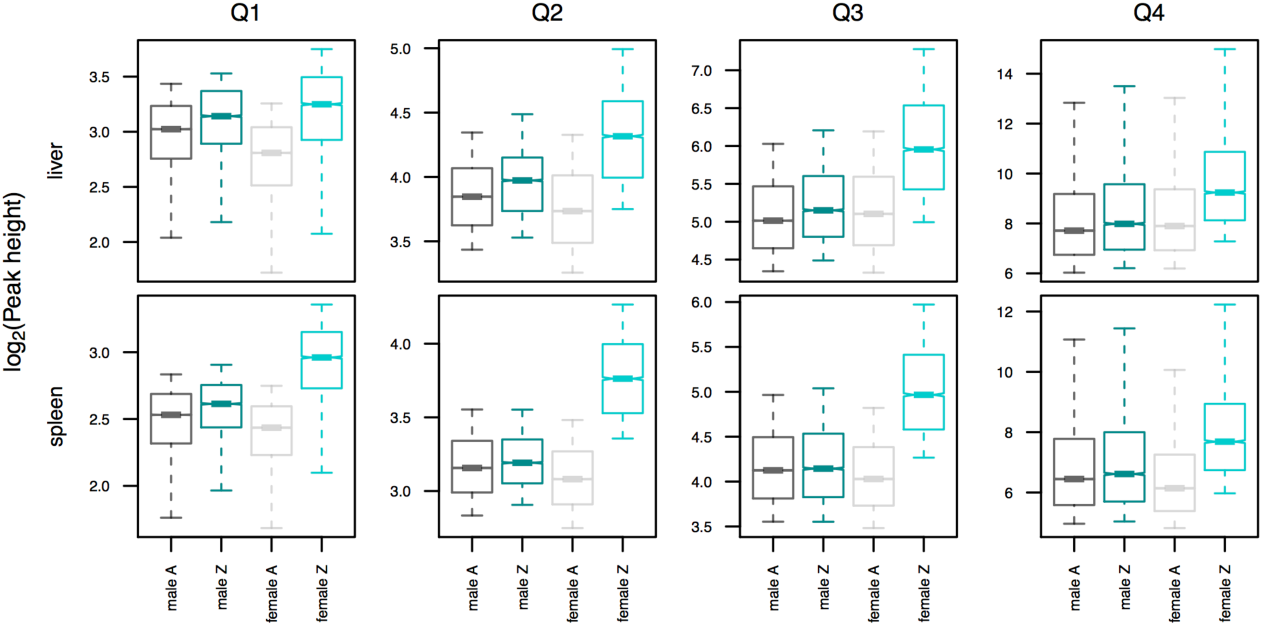
Boxplots presenting log_2_ gene-centred peak height distribution broken down into four quantiles (Q1 – Q4). A: autosomes, Z: Z chromosome. Upper panel liver, bottom panel spleen.

**Figure SI.**
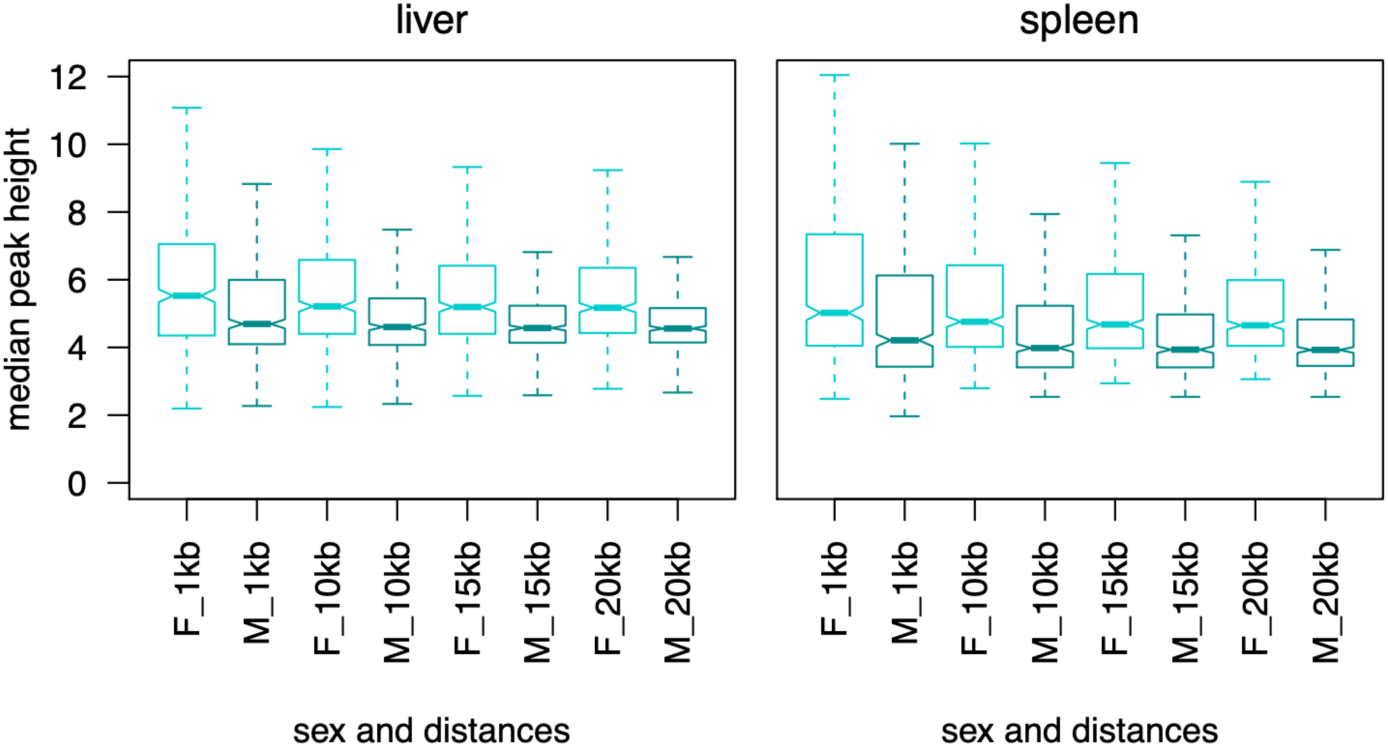
Boxplots showing gene-centred peak height distribution including differently sized regions around the gene body (1 kb, 10 kb, 15 kb and 20 kb) in females (F) and males (M) on the Z chromosome.

**Figure SJ.**
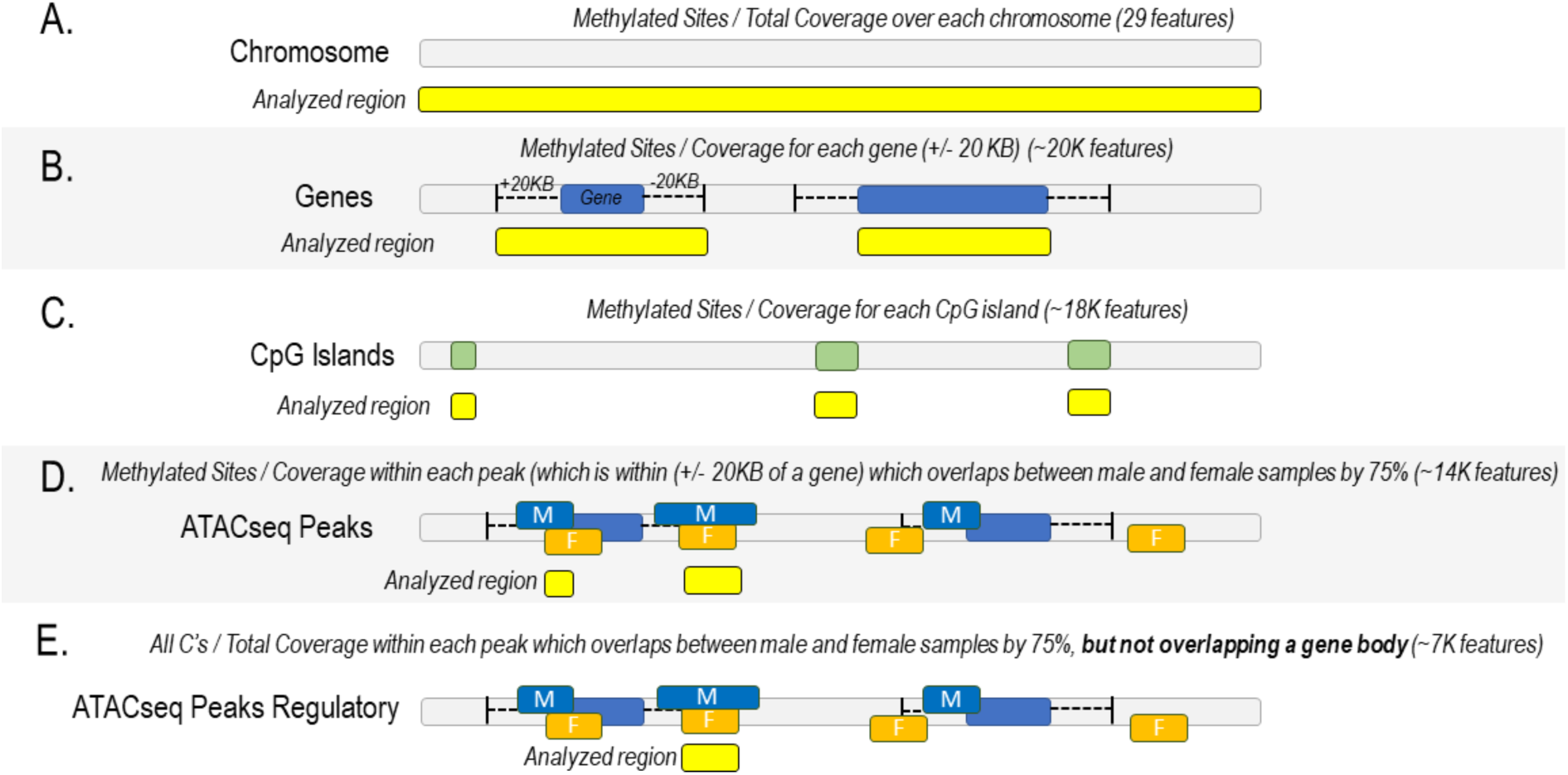
Cartoon depicting the five feature-based analyses used to examine 5mC DNA methylation. DNA methylation levels for each feature were calculated as the cumulative proportion of methylated reads (sum of methylated reads divided by the total number of reads per feature).

**Figure SK.**
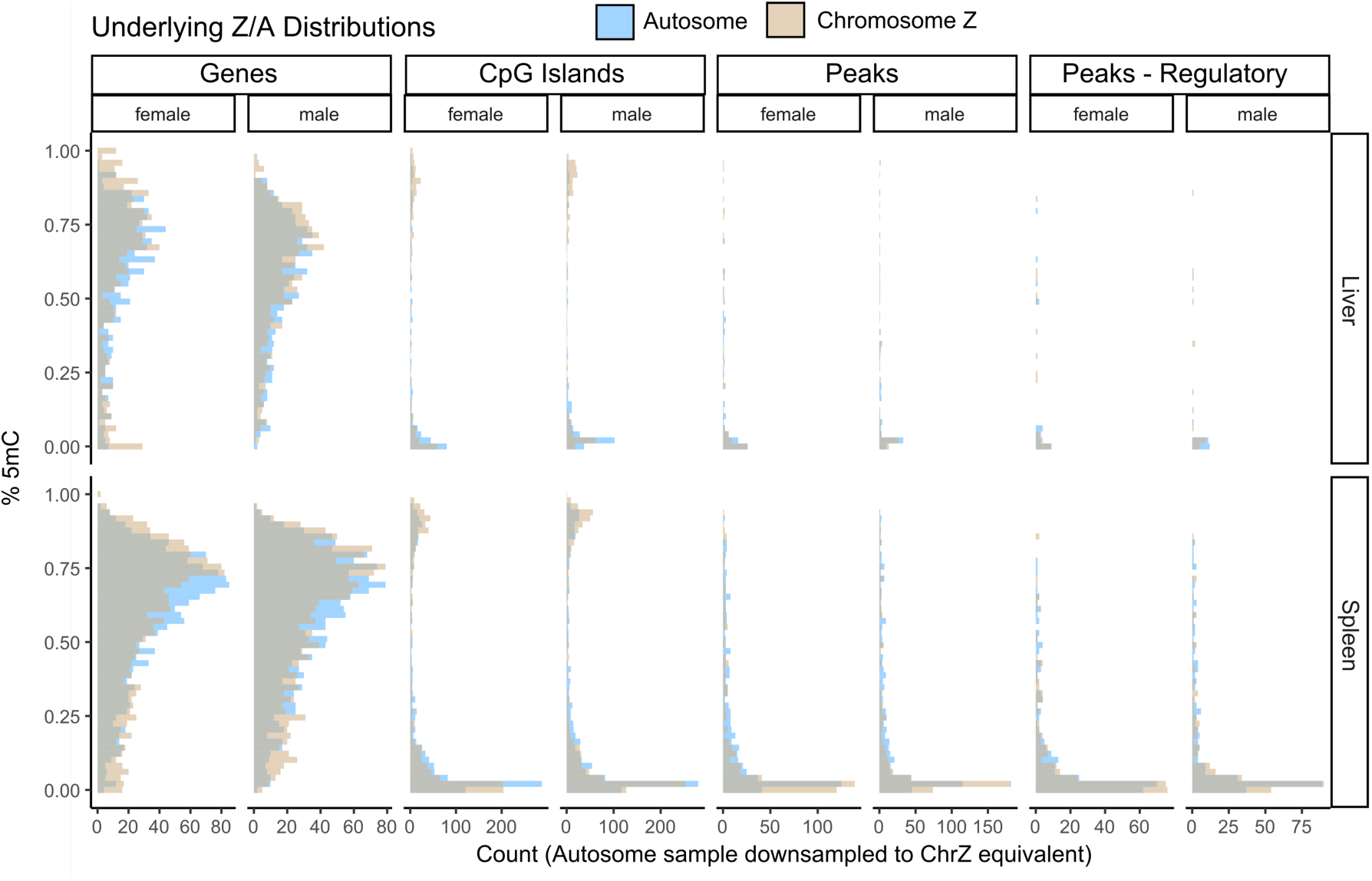
Frequency distributions of 5mC CpG DNA methylation underlying data in **Figure SL** and in **Supplementary Table A** for four of the feature categories (chromosome excluded because of a single feature unit for Chromosome Z). Autosomal distributions were randomly subsampled to be equivalent to the number of features present in the Z chromosome.

**Figure SL.**
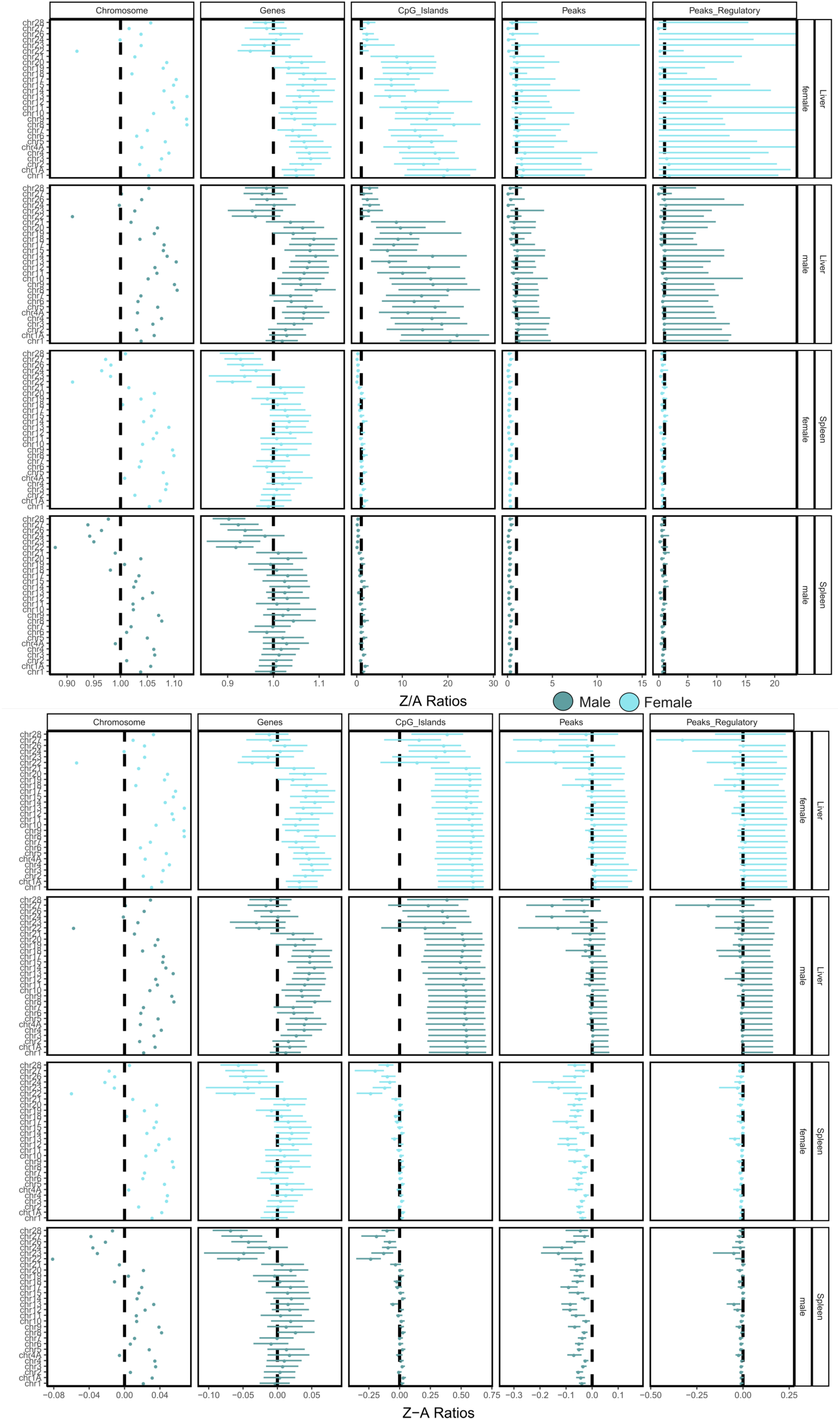
5mC CpG DNA methylation differences between chromosome Z and autosomal chromosomes for both sexes and tissues. Distributions were drawn from 10,000 bootstrap replicate samplings of the median. **A.** Ratio of DNA methylation on chromosome Z relative to the chromosome along the Y axis. Note variable X axes across facets. **B.** Same as panel A, except indicating the absolute difference in methylation (Chromosome Z – Chromosome N).

**Figure SM.**
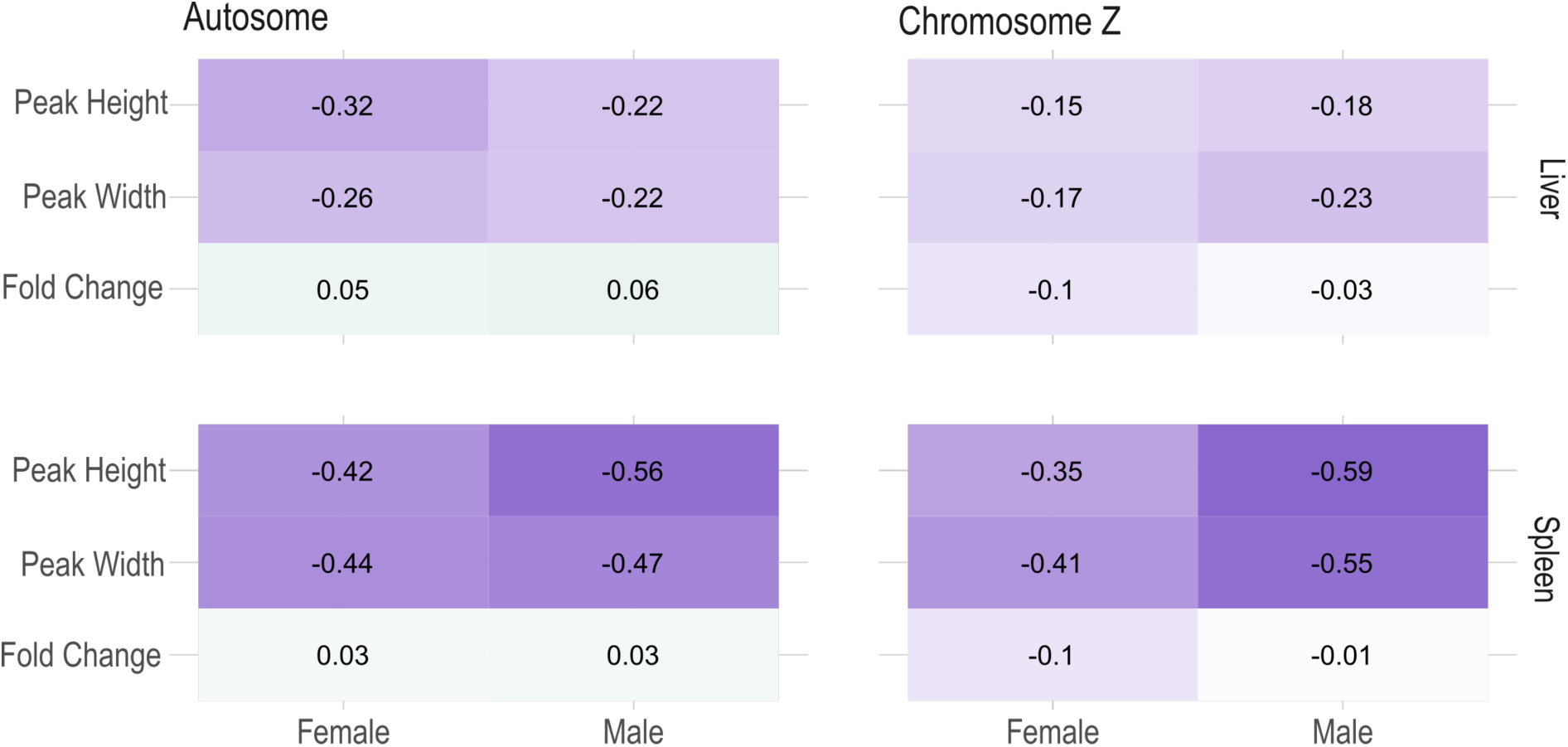
Spearman correlations between 5mC CpG methylation level and gene expression (FPKM) and open chromatin (peak height, peak width), calculated within each feature for each sex and tissue individually.

**Figure SN.**
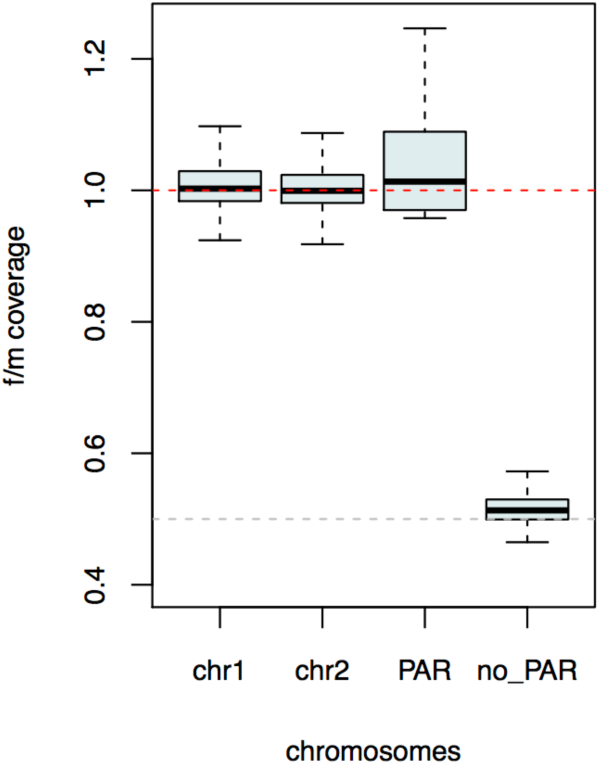
Boxplot of female to male read coverage distribution from whole genome shotgun sequencing of one individual of each sex as calculated in 50 kb windows, in chromosome 1 (chr1), chromosome 2 (chr2), the pseudoautosomal region (PAR) and the Z chromosome without the PAR. F/M coverage ratio on the shown autosomes and the PAR lie near a ratio of 1 (red line), whereas the hemizygous part of the Z lies close to a ratio of 0.5 (grey line).

**Figure SO.**
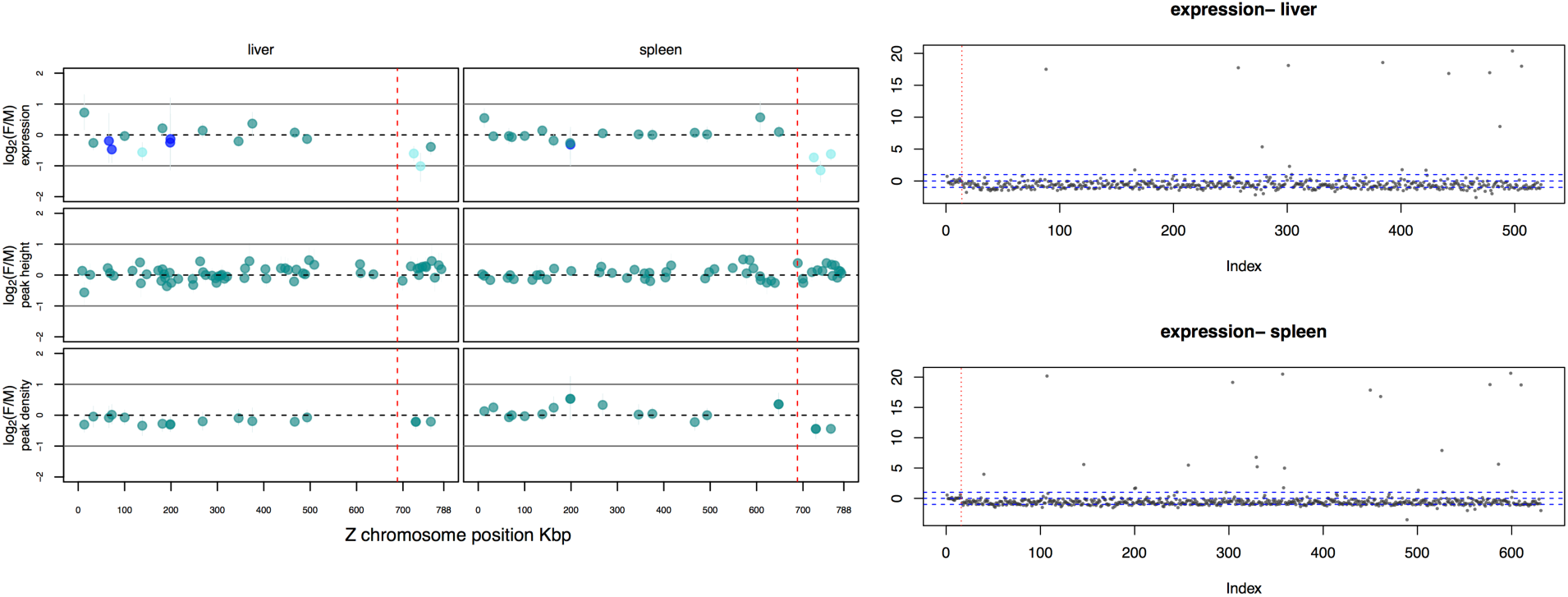
Left panel: The PAR region on the Z chromosome is shown in spleen and liver. Upper panel: log_2_(F/M) gene expression. Middle panel: log_2_(F/M) peak height. Bottom panel: log_2_(F/M) peak number. Right panel: log_2_(F/M) gene expression of all genes expressed in the Z. Red dotted vertical line marks the PAR.

**Figure SP.**
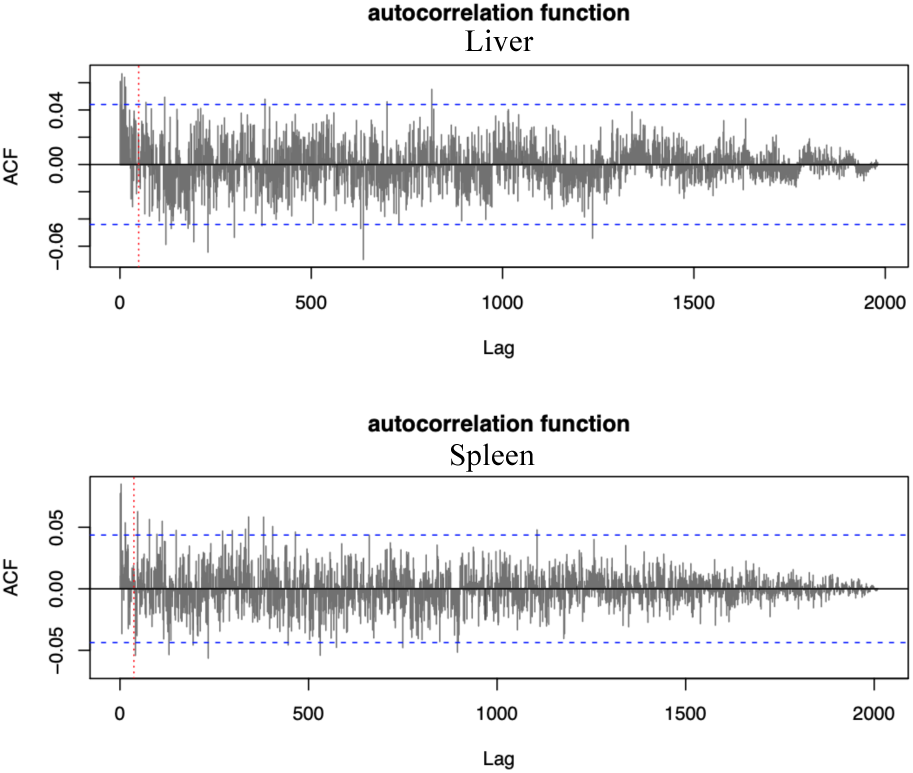
Autocorrelation of gene expression present on the Z chromosome. Red vertical line shows the PAR border. Blue lines represent autocorrelation values significantly different from zero. Except for the PAR region, there is no clear trend of clusters that would point at dosage local centers of dosage compensation.

**Figure SQ.**
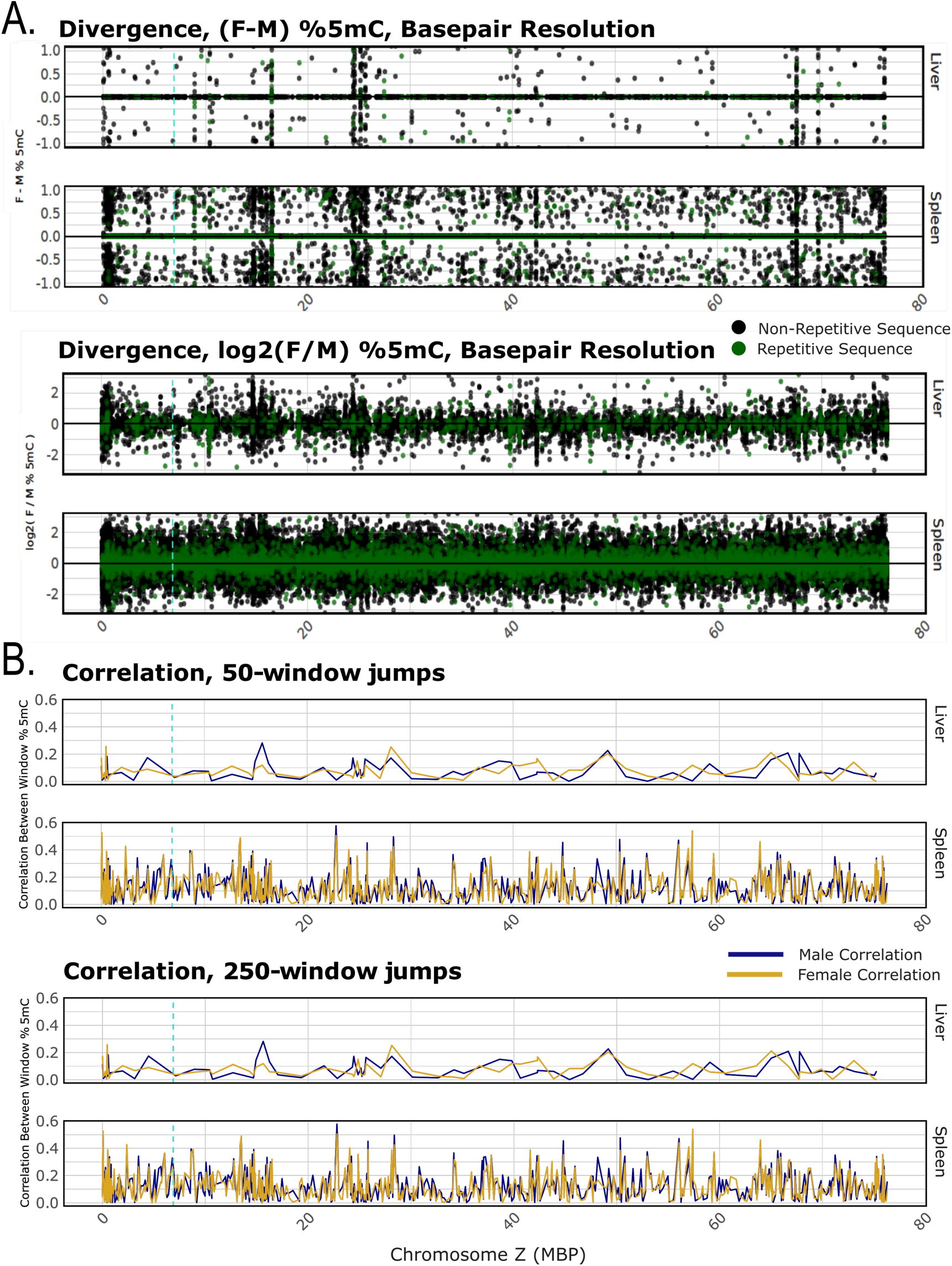
Sex-specific DNA methylation patterning along the Z chromosome at base-pair resolution calculated for both tissues and sexes. **A.** Divergence values shown for the difference in 5mC methylation between females and males (f-m; top) and the ratio between sexes (log2(f/m); bottom) along the Z chromosome suggest interspersed blocks of increased divergence, concomitant for both sexes. There was no evidence of a male-only hypermethylated region. **B.** Genomic autocorrelation between *n-*sized base-pair blocks. Spearman correlations were calculated between the first *n*/2 and last *n*/2 methylation calls in a block. Blocks were analyzed in both 50 and 250 base-pair blocks. We hypothesized that an MHM region in crow would exhibit male-specific hypermethylation, and strong male genomic autocorrelation.

**Figure SR.**
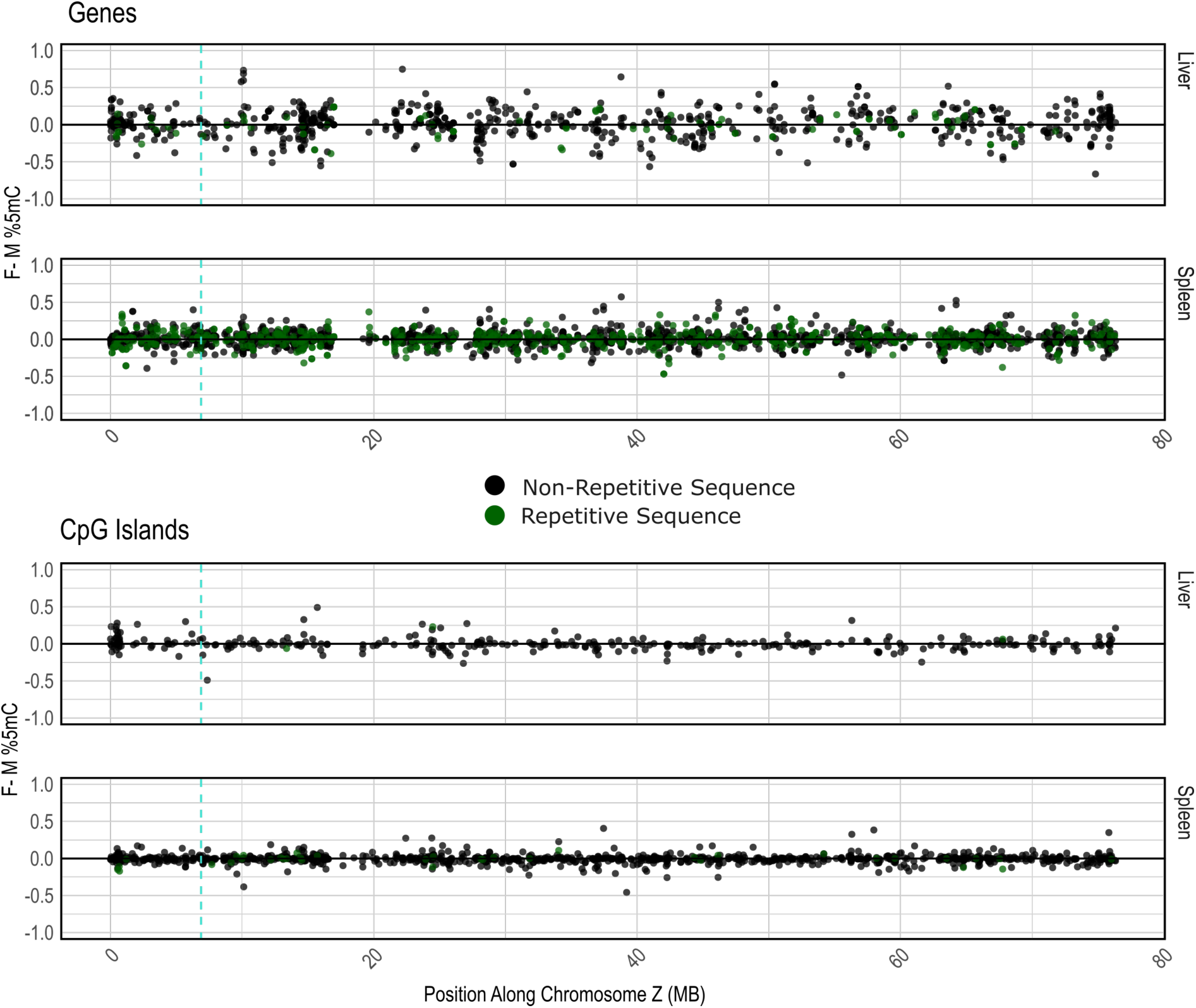
Sex-specific DNA methylation patterning along the Z chromosome shown as the difference between female and male methylation levels (f-m). Similar to Figure SQ, except instead of base-pair resolution, we examined DNA methylation divergence over genes and CpG islands.

**Figure SS.**
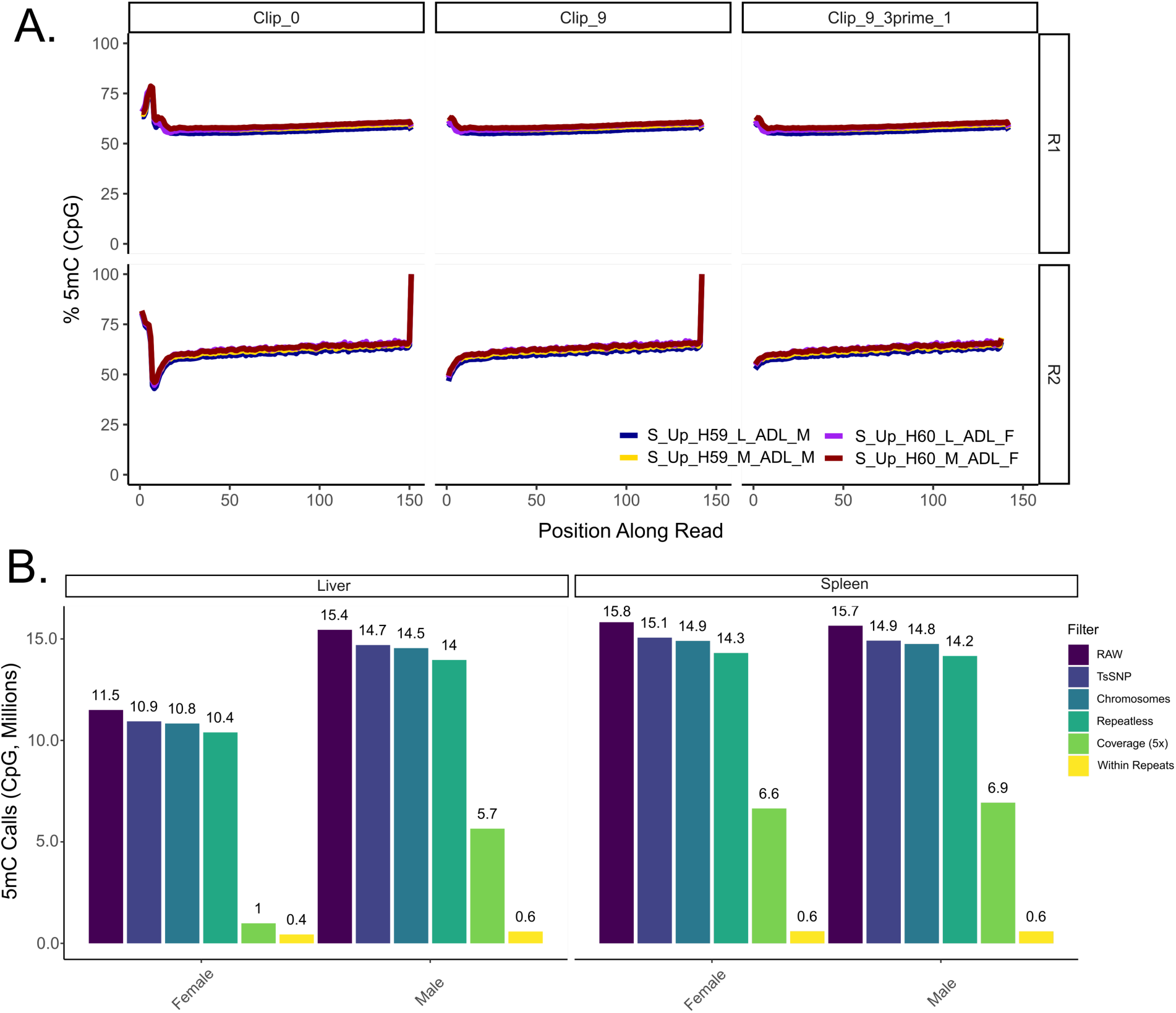
Results of quality control for whole-genome bisulfite-sequencing libraries for male and female hooded crow samples, created for both spleen and liver. Panel **A** indicates methylation levels averaged across all reads in a library along read position, used to identify systematic methylation biases along read lengths, with the far-right panel indicating our final cleaned reads. As our libraries were created with a post-bisulfite conversion kit, we trimmed the first 9 reads from the 5’ end, alleviating this known bias. We then iteratively revisited this step by trimming the last base from the 3’ end, due to the elevated 5mC levels in the last position on the R2 read. Sample name: Individual: S_Up_H59/H60; tissue:liver (L) or spleen (M); age: adult (ADL); sex: male (M) or females (F)). Panel **B** indicates total 5mC positions called for each sample, as well as sites retained after filtering for C-T and G-A SNPs, retaining sites on assigned chromosomes, removing sites within identified repeats, and filtering for 5x coverage.

